# Genomic insights into *Lactobacillaceae*: Analyzing the “Alleleome” of core pangenomes for enhanced understanding of strain diversity and revealing Phylogroup-specific unique variants

**DOI:** 10.1101/2023.09.22.558971

**Authors:** Archana S. Harke, Jonathan Josephs-Spauling, Omkar S. Mohite, Siddharth M. Chauhan, Omid Ardalani, Bernhard Palsson, Patrick V. Phaneuf

## Abstract

The *Lactobacillaceae* family’s significance in food and health, combined with available strain-specific genomes, enables genome assessment through pangenome analysis. The ‘Alleleome’ of the core pangenomes of *the Lactobacillaceae* family, which identifies natural sequence variations, was reconstructed from the amino acid and nucleotide sequences of the core genes across 2,447 strains of 26 species. It comprised 3.71 million amino acid variants in 29,448 core genes across the family. The alleleome analysis of the *Lactobacillaceae* family revealed key findings: 1) In the core pangenome, amino acid substitutions prevailed over rare insertions and deletions, 2) Purifying negative selection primarily influenced core gene variations in the family, with diversifying selection noted in *L. helveticus*. *L. plantarum*’s core alleleome was investigated due to its industrial importance. In *L. plantarum*, the defining characteristics of its core alleleome included: 1) It is highly conserved; 2) Among 235 isolation sources, the primary categories displaying variant prevalence were fermented food, feces, and unidentified sources; 3) It is predominantly characterized by ‘conservative’ and ‘moderately conservative’ mutations; and 4) Phylogroup-specific core variant gene analysis identified unique variants (DltX, FabZ1, Pts23B, CspP) in phylogroups ‘I’ and ‘B’ which could be used as identifier or validation markers of strain or phylogroup.

## 1. Introduction

The collective *Lactobacillaceae* family consists of bacteria with characteristic features of being gram-positive, acid-tolerant, fermentative microorganisms that produce lactic acid as the primary product in carbohydrate metabolism (George, 2018). The *Lactobacillaceae* family comprises 31 genera, 25 of which were formerly categorized under the genus *Lactobacillus* (Zheng et al., 2020). These microorganisms are naturally found across diverse niches including mucous membranes, mammalian gastrointestinal tracts, vagina, plants, fruits, vegetables, cereal grains, wine, milk, and meat environments; *Lactobacillaceae* are also historically applied as starter cultures in fermentation to produce dairy products (cheese, yogurt, dahi), meats, and vegetables, and also in making of wine, beer, coffee, silage, cocoa, and sourdough (Pasolli, 2020; Leech et al., 2020; Chee et al., 2020). The pivotal significance of *Lactobacillaceae* members in the realms of food, health, and probiotic industries, along with the enhanced affordability of sequencing technology, has ignited extensive research endeavors. As a consequence, a surge of *Lactobacillaceae* genome sequences has been generated, facilitating comprehensive large-scale genome analysis (Zheng et al., 2015). Many members of the *Lactobacillaceae* family are recognized by European Food Safety Authority (Bernardeau et al., 2008), with a few strains from the family marketed as probiotics (Klaenhammer et al., 2012). The probiotic products containing *Lactobacillaceae* have a substantial position in the probiotics market (Saxelin, 2008), with *L. plantarum, L. casei, L. helveticus, and L. acidophilus as* significant members contributing to and dominating the probiotics global market (Saxelin et al., 2005).

The pangenome represents the set of orthologous and unique genes present in a particular species, and the core genome of the pangenome represents the set of homologous genes shared by all analyzed strains (Costa et al., 2020). Sequence diversity studies across the *Lactobacillaceae* family and other closely related microorganisms enable insights on genetic similarity between strains through grouping classes by phylotyping (Silvaraju et al., 2021; Ellegaard et al., 2015; Spratt, 1999) based on the analysis of “Core” or “Accessory” genomes. It has led to the identification of crucial variations in genes (‘alleles’) relating to changes in pH, temperature, and salts or adaptations in response to niche requirements (Siezen et al., 2011; Chaillou et al., 2009) as well as to the identification of strains of probiotic potential concerning stress resistance (Arnold et al., 2018).

Intra-species sequence variation studies are essential for lactic acid bacteria as they highlight strain and species diversity, unveiling probiotic potential, functional abilities, and ecological adaptations. These insights have significant implications in healthcare, the food industry, and environmental research. Additionally, such studies enable the exploration of the genetic population structure of a particular organism. The concept of “ORF Alleleome”, where the “alleleome” signifies the aggregate of all gene alleles within an organism, provides a comprehensive understanding of genome-scale sequence variations, not only between the strains of a single species but also between different species within a family.

Here we report the ORF-alleleome analysis derived from the core pangenome of the *Lactobacillaceae* family at nucleotide and amino acid levels. This report explores the natural intra, and inter-species sequence variation patterns and hints at the links between the variants and their predicted substitution effects in this family. It also uncovers unique variants specific to phylogroups and strains, which could be used as markers to identify or validate strains. These sequence variations and unique variants could influence the functionality of the microbes in the Lactobacillaceae family. They could lead to differences in metabolism, stress tolerance, and probiotic effects, among other things. For example, some variations might enhance the ability of the bacteria to compete in the human gut environment, modulate the immune response in beneficial ways, or improve the flavor and texture of the food products. By examining these genetic differences in conjunction with experimental data on the bacteria’s function and behavior, it might be possible to draw connections between specific genetic variants and specific functional effects. These insights could guide the development of new probiotic strains with enhanced or tailored functionality.

## 2. Materials and methods

### 2.1 Pangenome procurement, sequence analysis, and quality control

The core genomes of all 26 species from the pangenome of *the Lactobacillaceae* family, were used for the current study. The number of core genes (Figure S1 A) and the number of strains varied for each species (Figure S1 C). The core genes (nucleic acid and amino acid sequence) with a single ORF were used for the core alleleome generation. The quality control and analysis methodology were adopted from (Catoiu et al., 2023) with certain modifications. The length of the nucleic acid sequence of each allele for each core gene was calculated. The nucleic acid alleles found to be more than two SDs shorter than the mean nucleic acid sequence length were removed using their ‘Locus id,’ and a similar set of alleles was also removed from the amino acid alleles (Figure S1 B). The alleleome of codon sequence variation and amino acid sequence variation was generated using the final nucleic acid and amino acid alleles, respectively (Figure S1 B).

### 2.2 Alleleome analysis workflow and core alleleome generation

The alleleome was defined by following a strategic workflow (Figure 1, explained with *L. plantarum* as an example species). This strategy was used to define the core alleleome of 26 species of the *Lactobacillaceae* family. Multiple sequence alignment (MSA) of alleles of each core gene was performed, followed by the consensus sequence construction. The consensus sequence was derived using Biopython (Chapman and Chang, 2000) with a threshold frequency of 0.5 (50%) by obtaining the amino acid present in above 50% of the strains to acquire the amino acid with a dominant presence. MSA was performed using MAFFT (Multiple sequence Alignment using Fast Fourier Transform) tool (Katoh and Toh, 2008). Each allele sequence was aligned with the consensus sequence using stand-alone BLAST (Madden, 2003) to obtain the variants. The variants of type substitution, insertion, and deletion were determined. The in-house designed and developed Python (Van Rossum and Drake, 2009) scripts were used for the core alleleome generation.

**Figure 1:**
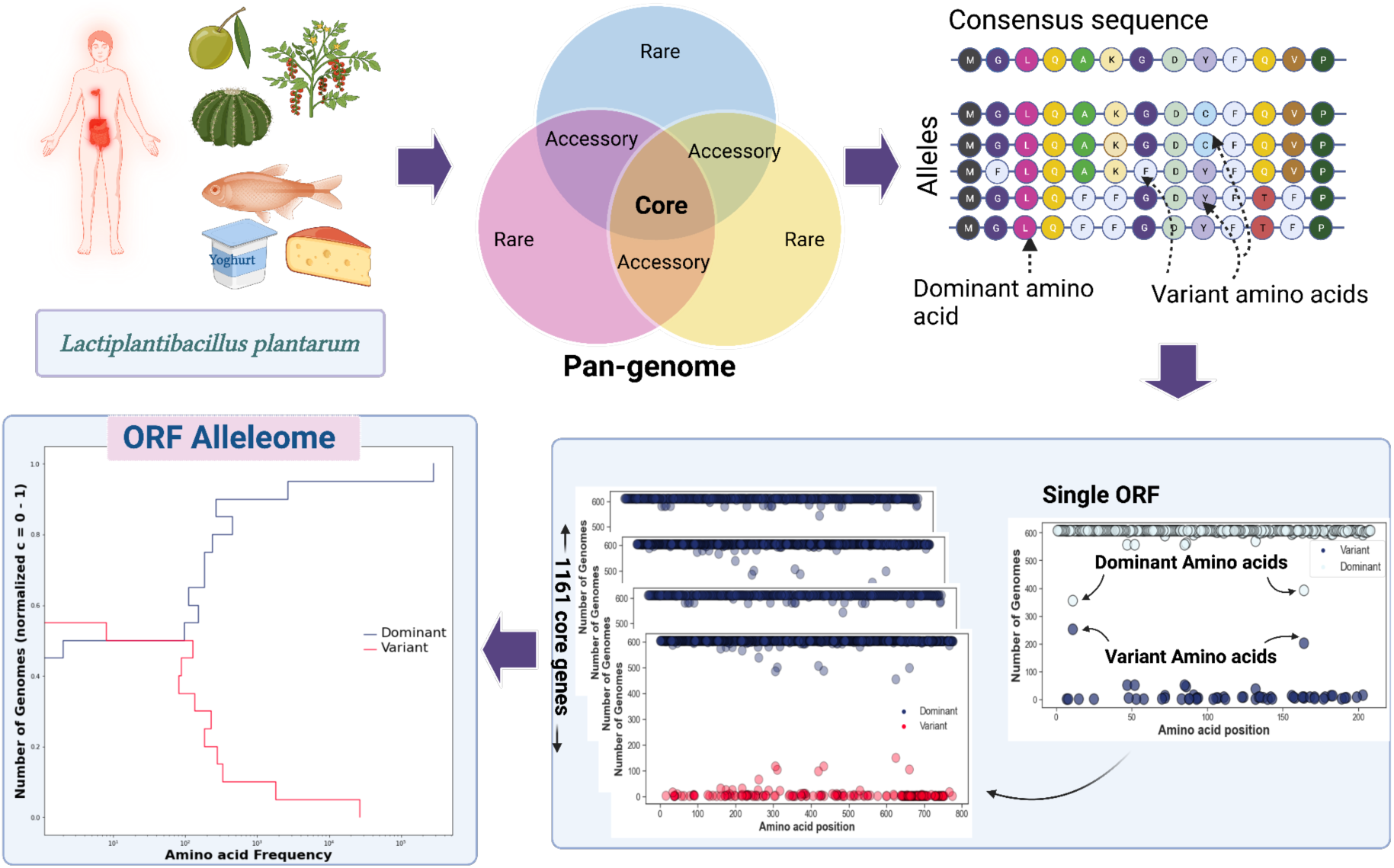
Alleleome workflow describing the strategy of alleleome generation.

### 2.3 Determination of variant distribution across core pangenome

The percentage distribution of variants (substitution, deletion, and insertion) within the core genes of the *Lactobacillaceae* family was analyzed from different viewpoints to provide insights into the overall mutation patterns within these species (Figure 2). The count of each mutation type and their corresponding percentage were calculated across the core pangenome of each species (Figure 2A). The correlation between the variants count, and the number of core variant genes for each strain was calculated to understand the potential interplay between core genes and variability (Figure 2B). The unique variants per core variant gene per species were calculated by considering every distinct substitution in the consensus sequence and the variants (Figure 2C), followed by further calculating the average of those unique variants (Figure 2D).

**Figure 2:**
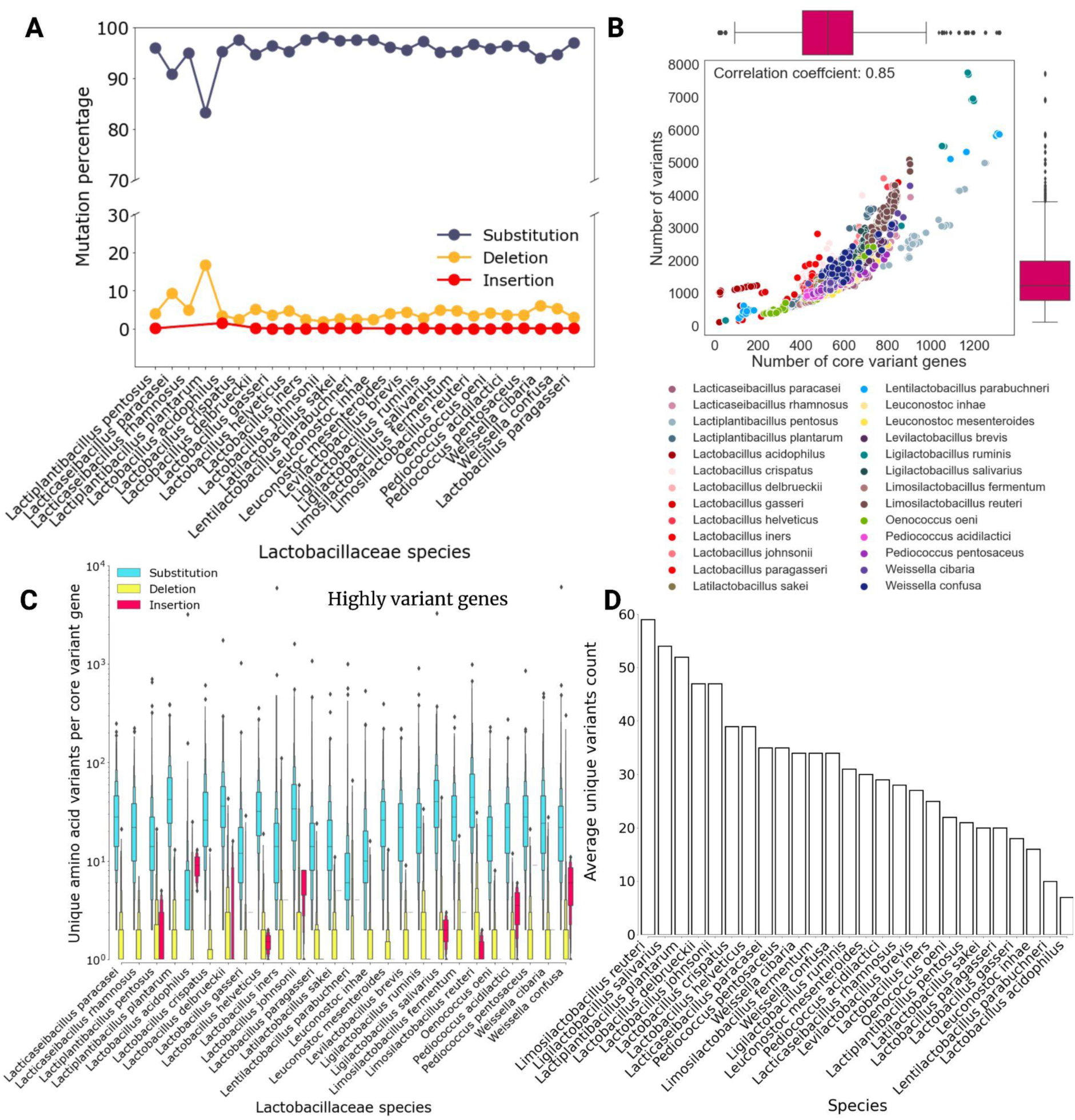
A) Distribution of variants across core pangenome of all 26 species; B) Correlation of the number of core variant genes and number of variants in each strain in all 26 species; C) Distribution of unique amino acid variants per core variant gene with mutation type (substitution, insertion, and deletion) in all species; D) Average number of unique variants per core variant gene in all 26 species ranked from higher to lower values.

### 2.4 Determination of mutation rates of codon changes and identification of selection pressures

The methodology in (Catoiu et al., 2023) was adapted and implemented to determine the mutation rates of codon-specific changes for all 26 species core alleleome. In the alleleome, the dominant codon (‘Consensus codon’) and non-dominant codon (‘Variant codon’) at the same position were determined through the consensus sequence generation and the alignment of each allele with this consensus. The total number of strains represented at the position by the ‘Variant codon’ was counted as the number of mutations from the consensus sequence. Next, the count of mutations with the specific codon change (‘Consensus codon’ → ‘Variant codon’ pair) was pooled to determine the rates of specific codon changes across the alleleome. These codon changes were further separated by their effect on the amino acid (synonymous vs. nonsynonymous amino acid substitution). The rates of synonymous and nonsynonymous mutations (dN/dS) per core gene were calculated and used to extrapolate the general selection pressures acting on each gene in each species. These ratios were calculated for the alleleome of all 26 species. These values were plotted for core genes mutated in the strains at the family level and species level (Figure 3A, 3B) and the bar plot of the count of core genes with dN/dS ratio in three categories (dN/dS < 1, dN/dS > 1, dN/dS = 0) was also plotted for individual species (Figure 3C).

**Figure 3:**
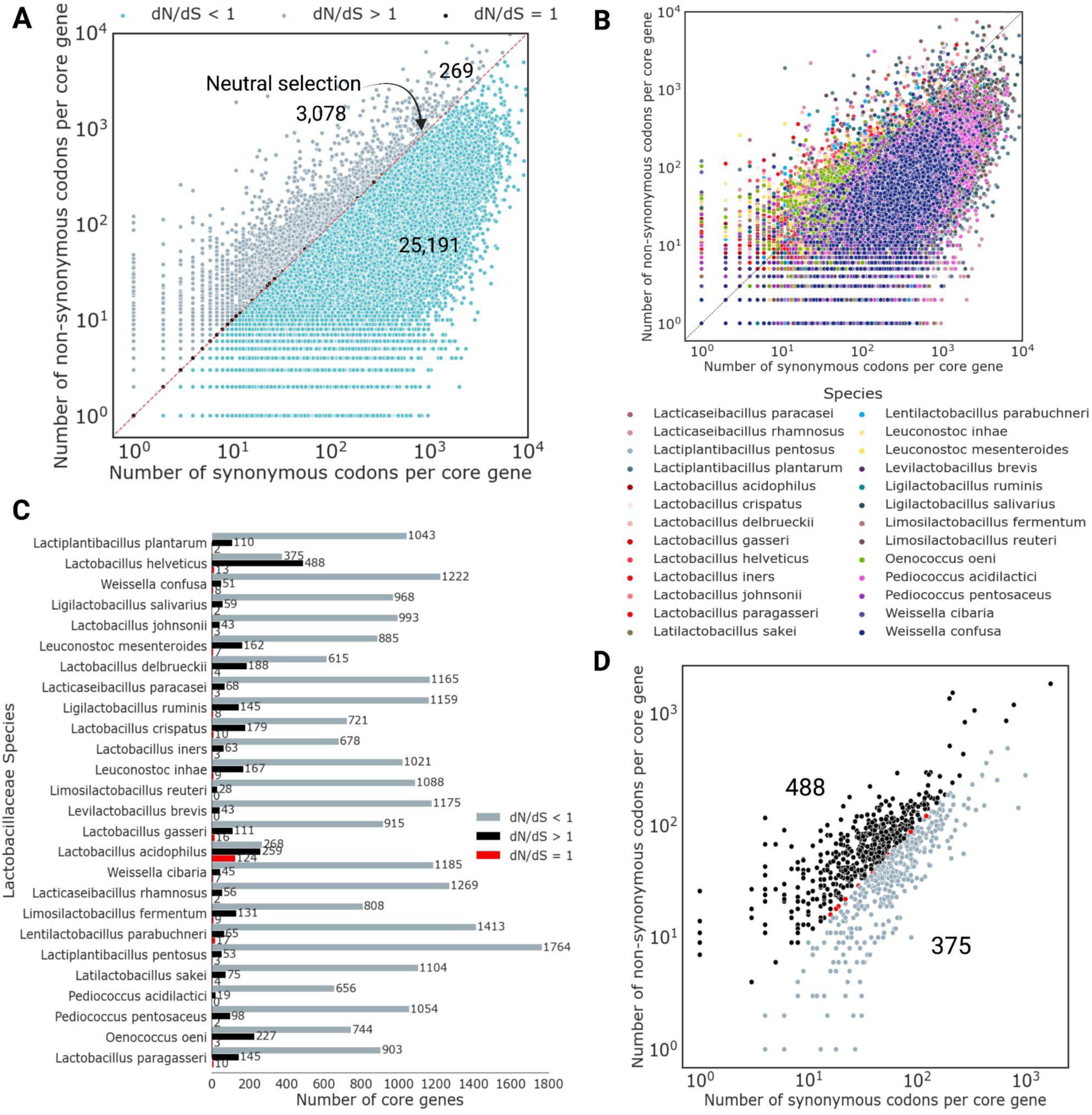
A) The number of synonymous and nonsynonymous mutations in each gene in Lactobacillaceae family where genes are colored by their dominant selection pressure: purifying selection (cyan), diversifying selection (grey), and neutral selection (black); B) The number of synonymous and nonsynonymous mutations in each core gene in Lactobacillaceae family colored by 26 species; C) Number of core genes with dN/dS ratio values categorized in three levels (dN/dS > 1, dN/dS = 1, dN/dS < 1) in each species; D) L. helveticus with diversifying positive selection pressure.

### 2.5 Principal Component Analysis (PCA) of amino acid substitutions, COG categories, and Grantham scores

First, the COG categories from pangenome analysis (Rajput et al., 2023) were mapped with the core variant genes in the core alleleome of all 26 species, and the percentage distribution of all variant types (substitution, insertion, and deletion) across all COG categories was calculated. Their distribution was plotted using a heatmap (Figure S2).

Further, a PCA methodology was utilized to elucidate patterns and assess if a correlation exists between the amino acid substitutions (consensus and variant amino acid), COG categories, and predicted substitution effects across 26 species (Figure S3-1). The features included for PCA were the unique amino acid substitution count (substitution count) across the core alleleome of each species, consensus amino acid, variant amino acid, COG category, Grantham score, and species name. For the substitution effects, the Grantham scores were calculated for each amino acid substitution. PCA enables the reduction of the dataset into fewer dimensions while retaining key trends and patterns, thus simplifying data visualization and interpretation. Although this package is specifically designed to handle mixed data, it can also perform the PCA of only quantitative or qualitative variables (Chavent et al., 2022). PCAmixdata performs this dimensionality reduction using the singular values decomposition method (Chavent et al., 2022).

A PCA was performed on the dataset, which included amino acid substitutions (consensus and variant), COG categories, substitution count, and Grantham scores across all the species. All the features except substitution count and Grantham score were categorical variables. The categorical variables were converted to numerical data by the distribution of the numbers/fractions between 0 and 1 based on the unique values of each feature. For example, the consensus and variant amino acids had 20 unique amino acids. Hence, the numerical conversion for these features included a distribution of 0 to 1 in 20 numbers/fractions. Likewise, this conversion method was applied to all the categorical features. The eigenvalues, the percentage of variance, the cumulative percentage of variance (Table S1), and square loadings (Figure S3-1A) for six PCs were calculated. The distribution of observations across the first two PCs was colored by important features (Figures S3-1B, S3-1C, S3-1D, S3-2A, S3-2 B, S3-2C).

### 2.6 Construction of *L. plantarum* core alleleome

The core alleleome was constructed using the methodology of (Catoiu et al., 2023) with trivial modifications. The occurrence of dominant and variant amino acids was calculated to obtain their count across the core alleleome at each amino acid position in the MSA alignment. The amino acid variant count was obtained based on its presence in the number of strains, and the dominant amino acid was the amino acid of highest occurrence obtained in the consensus sequence at each position. When the dominant amino acid is a “deletion” (for instance, when an insertion is detected in strains), that residue position is excluded from the overall consensus sequence.

The in-frame deletions and insertions (for those species where insertions were observed) were counted as amino acid variants in the core alleleome (Figure 4). However, those were not considered in the Gratham score analyses (Figure 5), as the Grantham score is calculated based on the amino acid combinations in the substitutions.

**Figure 4:**
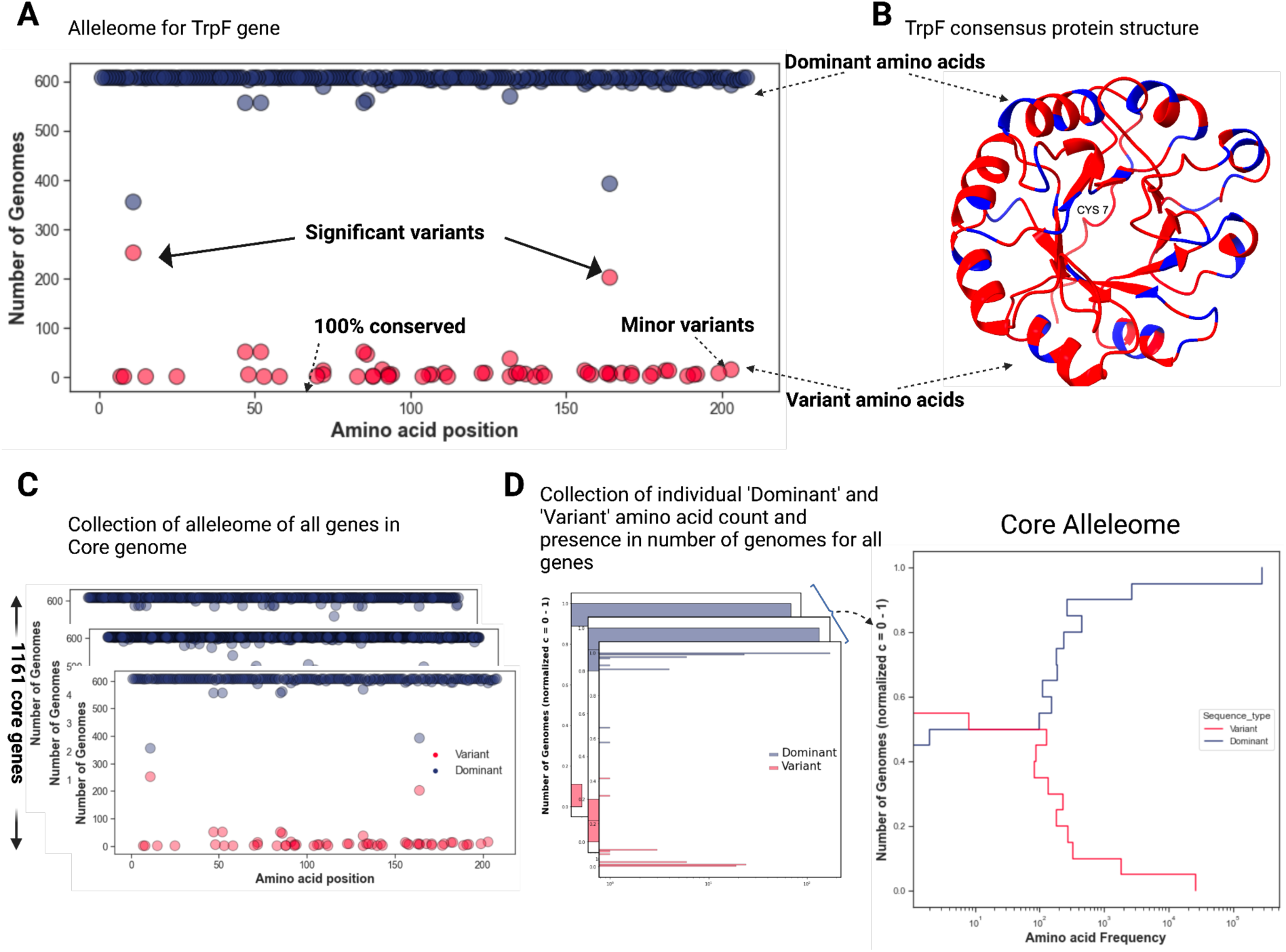
A) ‘Dominant’ (of highest occurrence) and ‘Variant’ (non-dominant) amino acids and their respective natural occurrences for amino acid positions; B) The substitutions mapped onto the predicted consensus structure; C) Collection of all the ‘Dominant’ and ‘Variant’ amino acids and their respective natural occurrences in all amino acid positions; D) L. plantarum core alleleome: Collection of all ‘Dominant’ and ‘Variant’ frequency without considering the positions. Conserved regions of the core alleleome are defined by positions where the dominant amino acid frequency is at least 90% (c ≥ 90).

**Figure 5:**
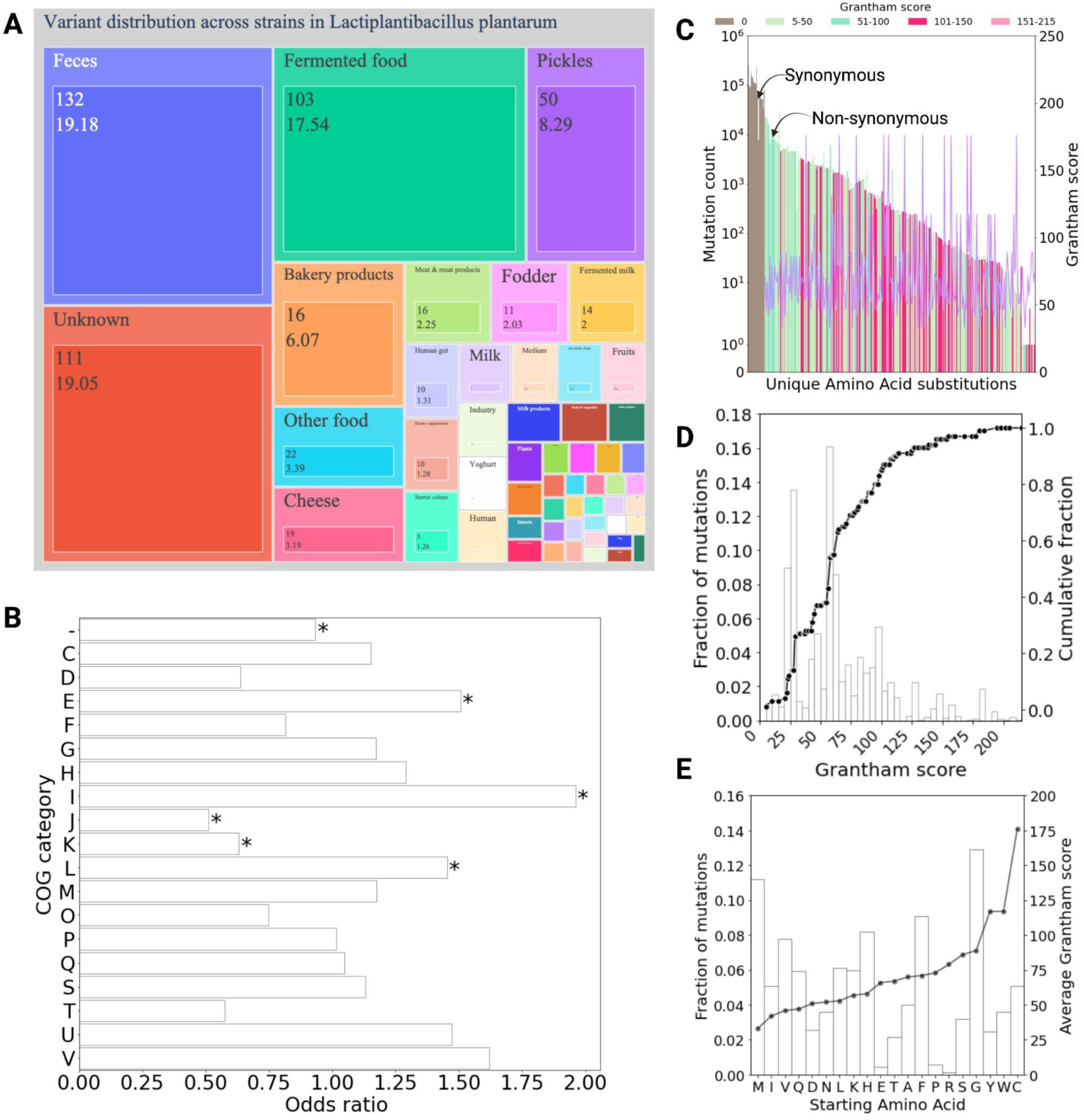
A) Strains grouped according to their isolation sources and colored according to the number of variants for L. plantarum. The size of the square represents the number of variants (in percentage) and the number of strains of the respective isolation source. B) Odds ratio from Fisher’s exact test of variants in COG categories and number of core genes across the alleleome; C) The alleleome of codon substitutions in terms of synonymous and non-synonymous substitutions with purple line showing the cumulative average of Grantham scores of non-synonymous substitutions; D) The Grantham score distribution across non-synonymous substitutions and black line showing their cumulative fraction of mutations; E) The nonsynonymous amino acid substitutions distribution with starting amino acid and gray line showing their average Grantham scores.

### 2.7 Distribution of variants across isolation sources and COG categories in core pangenome of *L. plantarum*

The 239 isolation sources were first segregated into 53 classes based on the source type. For example, ‘Kimchi’, ‘Okpei-Nsukka’, and ‘kinema’ were assigned to a ‘Fermented food’ class based on their preparation process. The percentage of variants and their occurrence in the number of strains was calculated for each class. This distribution explains the niches’ origin, strain specificity, and variant contribution (Figure 5A).

The count and percentage of variants assigned per protein-encoding core gene were calculated for all the unique (19) and combined COG categories (44). The COG variation distribution represents the extent of amino acid mutations within the life maintenance systems of an organism. With this point of view, only the key categories (19) were evaluated (Figure 5B).

Further, Fischer’s exact test was conducted to determine the enrichment relationship between the count of core variant genes where variants in COGs occurred and the number of variants present within each COG functional category. For instance, for COG category E, the test compared both COG E versus non-COG E core variant genes and COG E versus non-COG E variants. This procedure was replicated for 19 COGs, resulting in 19 separate tests. The significance threshold was set with a Familywise Error Rate (FWER) of less than 0.05, following a Bonferroni correction. Additionally, odds ratios were calculated as part of the statistical assessment.

### 2.8 Average Grantham score calculation and statistical analysis of Grantham score distributions in strains of *L. plantarum*

Each explicit codon mutation pair (‘Consensus codon’ → ‘Variant codon’) was counted across strains and further aggregated based on their mutation type (synonymous and non-synonymous) (Figure 5C). The non-terminating codons’ rates of synonymous and non-synonymous mutations were determined (61/64). Next, the distribution of Grantham scores was determined using the mutation rates for all non-synonymous mutations in strains (Figure 5D).

Further, the Grantham score of each mutation (amino acid substitution) and its frequency of occurrence in the non-synonymous core alleleome were used to find the average Grantham score of all amino acid substitutions originating from the same codon. This Grantham score represents the average severity of all mutations stemming from the same initial codon across all positions in the alleleome. The observed codon changes were also grouped by their premutation amino acid (Figure 5E). For all 20 amino acids, the distribution of all non-synonymous amino acid substitutions that originate from the same consensus amino acid is used to calculate the average Grantham score observed in a particular amino acid (Figure 5E). This Grantham score represents the average severity of all mutations stemming from the same initial AA across all positions in the alleleome.

The mutations are grouped by consensus amino acid (as discussed above), and the cumulative fraction of these Grantham scores was also determined. The cumulative average of the Grantham score (Figure 5C line), the cumulative fraction of mutations (Figure 5D black line), and the average Grantham scores were also observed (Figure 5E grey line).

### 2.9 Phylogenetic analysis of core genomes of *L. plantarum*

The core genes of 611 strains of *L. plantarum* were aligned using Roary (Page et al., 2015) and the aligned core genes were used for the construction of a phylogenetic tree. The aligned core gene sequences were first subjected to the ModelFinder Plus tool of IQ-TREE (version 1.6.12) to evaluate the best substitution model and further calculate the strains’ evolutionary differences. The phylogenetic tree was constructed using the best substitution model GTR+F+R10. The tree branches indicate the evolutionary differences in the core gene sequences between samples (Figure 6A). To further determine the association of these differences with the phylogroups and isolation sources, the strains were mapped with their phylogroups and sources, and the iTOL12 was used for presentation.

**Figure 6:**
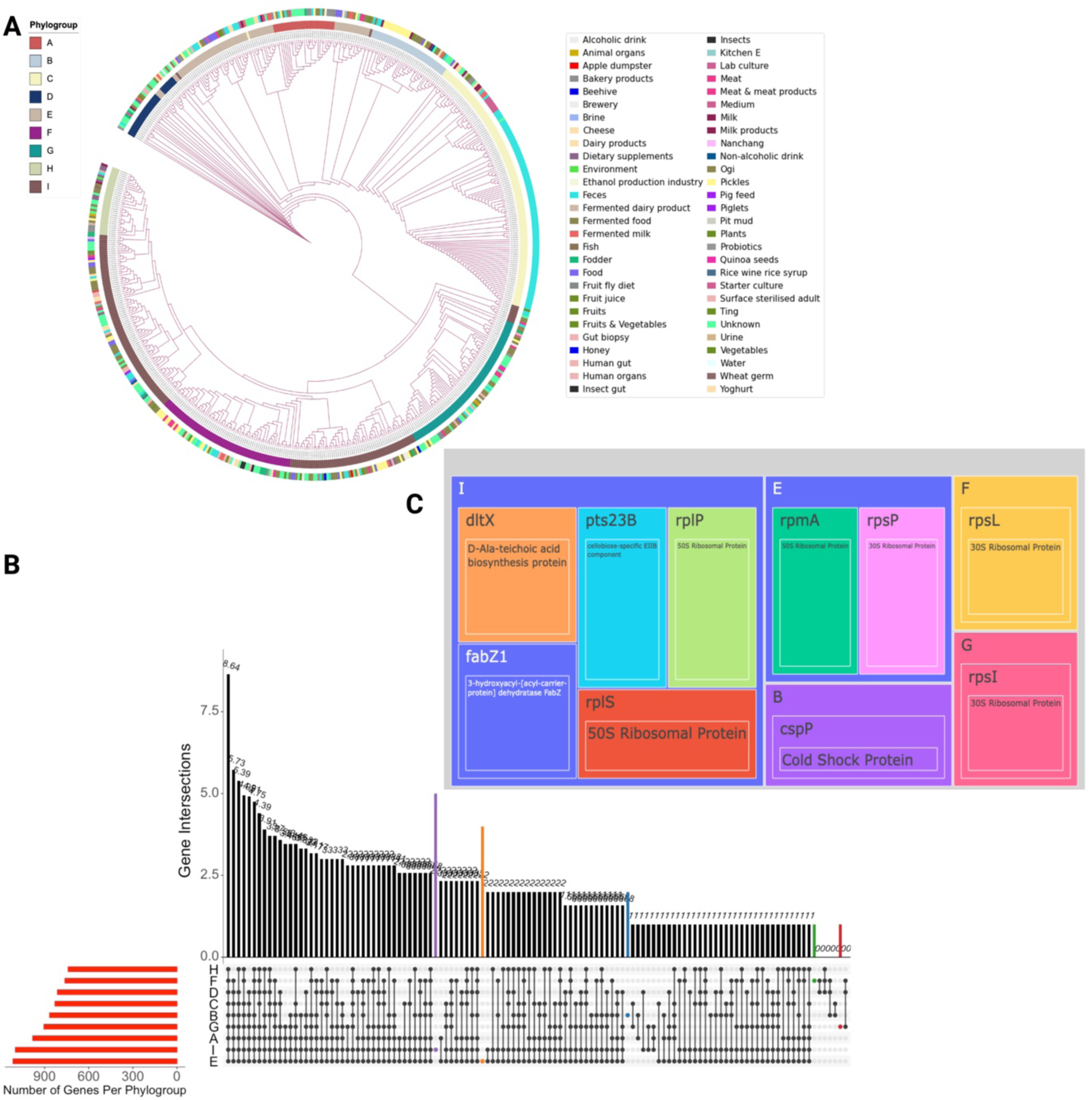
A) Phylogenetic analysis of core genomes of L. plantarum mapped with phylogroup and Isolation sources; B) Upset plot showing the intersections of core variant genes shared by phylogroups. The sidebar shows the total number of core variant genes per phylogroup. The intersection identifies the set of core variant genes shared by the phylogroups. The edges connecting nodes corresponding to the phylogroups signify their intersections. The red-colored bars correspond to the total count of variant core genes specific to each phylogroup. The black-colored bars signify the number of genes common to the respective groupings of phylogroups. Individual nodes marked devoid of connecting edges signify the core variant genes exclusive to specific phylogroups; C) The unique core variant genes and their proteins in specific phylogroups. The treemap displays the genes and their related functional annotations, with each phylogroup represented by a separate square. The specific genes and functional annotations of each phylogroup are denoted within these squares, and the coloration of the squares is determined by the genes.

### 2.10 Determination of Phylogroup-specific unique variants

The presence/absence matrix of the core variant genes associated with variants was constructed. The presence/absence matrix was a binary matrix, where the presence is denoted by one and the absence by zero. Further, the intersection operation was carried out on the elements shared by the phylogroups. A Venn diagram of the core variant genes associated with these variants and their phylogroups was constructed using intersection with UpsetR (Figure 6B). These intersections are represented by the edges connecting the nodes associated with the phylogroups. The phylogroup-specific unique core variant genes were identified (Figure 6C).

Further, the variants of these unique core variant genes were evaluated for their specific amino acid mutation sites and isolation sources (Figures 7A, 7B, and Figures S8A, S8B, S8C). The distribution of the unique variants across the alleles, along with the positions of the variants and strains, were represented by isolation sources and were colored by their predicted substitution effects. The consensus sequence of each phylogroup-specific unique core variant gene was used to construct the consensus protein structure. Alphafold2 (Jumper et al., 2021) using MMseqs2 was used on ColabFold (v1.5.2) (Mirdita et al., 2022) for the prediction of all the consensus protein structures. The variants were mapped onto the 3D consensus protein structures indicated by the variant position and the consensus amino acid at that position using Chimera X (Pettersen et al., 2021).

**Figure 7:**
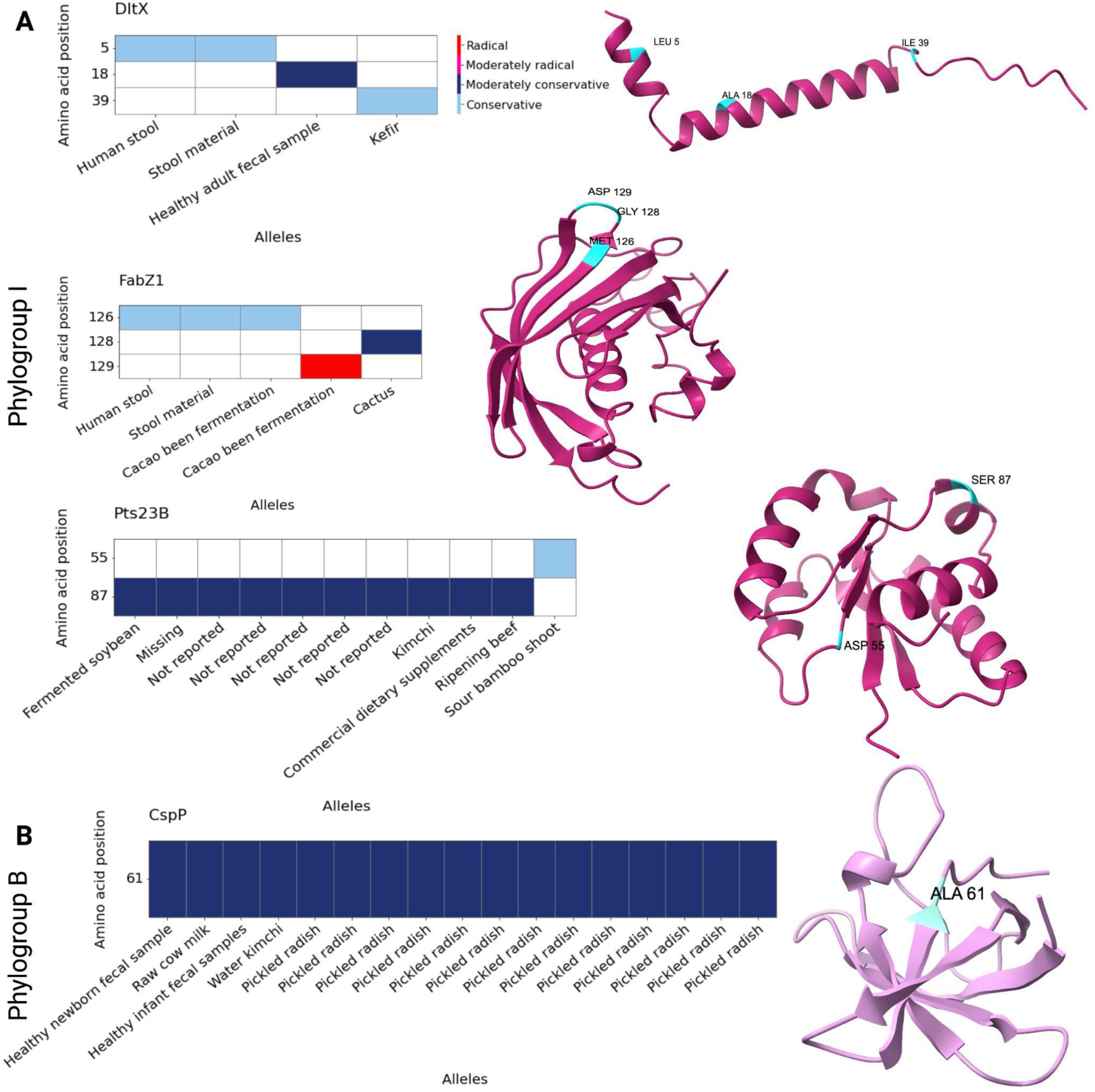
A) Phylogroup I specific variants of genes “DltX”, “FabZ1”, “Pts23B” with their amino acid mutation position, predicted substitution effects, and isolation sources; B) Phylogroup B specific variants of gene “CspP” with their amino acid mutation position, substitution effects, and isolation sources. The mutation positions are colored by their Grantham scores predicted effects for the genes.

## 3. Results

### 3.1 Core gene variability is not related to strain count and variant abundance

The study aimed to explore the natural amino acid sequence diversity within species. For this purpose, the variants in the core genes of the pangenome of 26 *Lactobacillus* species were analyzed. The core genome, consisting of 31,231 genes, was derived from 2,447 strains of the 26 *Lactobacillus* species. The amino acid variants were identified using a consensus-based approach (Figure 1), allowing for a more profound representation of the sequence variation within each species. This natural intra-species sequence diversity collection in protein-encoding core genes is referred to as core “ORF alleleome” (alleleome).

The core pangenome sequence diversity is a crucial representation of genes with essential functions to regulate biological processes. Hence the impact of variations on the functionality of core genes would be of interest. The alleleome of the core genes was reconstructed to interpret both the natural nucleotide and amino acid sequence diversity across 26 species. The analysis of natural sequence diversity revealed that among the 31,231 core genes in all *Lactobacillus* species, a majority of 29,448 core genes (94%) exhibited variations, including substitutions, insertions, and deletions. Over 90% of core genes showed variants in most species, except *L. acidophilus* (67%) and *L. parabuchneri* (82%). *L. acidophilus* showed the lowest variation percentage (67%, 1003 core / 1486 total genes), while *L. plantarum* exhibited the highest (99.66%, 1157 core / 1161 total genes) (Figure S1 A).

The *Lactobacillaceae* family members were further scrutinized to assess how strain or variant counts might affect the prevalence of core variant genes (Figure S1C, Figure 2A). To verify it, the correlation between them was examined. The correlation coefficient of core variant genes with the number of strains and variants count was 0.01 and 0.11, respectively (Figure S1 D). This demonstrated that the core variant gene counts are not correlated with the number of strains or variant abundance of a species. To illustrate, the species *L. gasseri* (31 strains), *L. helveticus* (31 strains), *L. parabuchneri* (30 strains), and *L. iners* (30 strains) with the lower number of strains had ‘34,221’, ‘66,009’, ‘69,570’ and ‘42,745’ variants, respectively (Figure 2A). Nevertheless, the percentage of core variant genes in these species was 94%, 99.5%, 82%, and 90%, respectively. Further, as stated above, the species *L. acidophilus*, with the lowest core variant genes, had 54 strains with 56,697 variants (Figure S1C). It is also to be noted that each strain had a different number of genes. Taken together, these results suggest that core genetic variation is not directly proportional to the number of strains or variants within a species. These findings also indicate that the percentage of core genes undergoing variations remains high regardless of the number of strains or variants in a species. Moreover, the discrepancy in the percentage of core variant genes is not related to the number of strains or variants in a species.

The initial total number of alleles of core genomes of all species included 2,887,790 before quality control and analysis (See method section 4.1), and after removing the sequences with length less than two standard deviation (SD) of mean length resulted in 2,868,484 alleles, with a loss of 19,306 (0.67%) (Figure S1 B).

### 3.2 The *Lactobacillaceae* family core pangenome is predominated with abundant substitutions and very low insertions and deletions (indels)

The core genome constitutes all the genes that are crucial for the fundamental biological functions and the major phenotypic features of a species. Variants within these genes could differentiate their functional capability, thereby affecting key aspects of the organism such as survival, metabolic competence, stress tolerance, probiotic characteristics (Lebeer et al., 2008), and organismal fitness (Kronenberg et al., 2018). To reveal these variants in the *Lactobacillaceae* family, the distribution of variations within the core genes, including substitutions, insertions, and deletions, was assessed by calculating their percentage for each species (Figure 2A). The study uncovered surprising findings, highlighting an abundance of substitutions, a lower percentage of deletions, and either a minimal presence or a complete absence of insertions in the *Lactobacillaceae* family.

The species with no insertions in the core pangenome were observed as *L. inhae, L. crispatus, L. plantarum, L. paracasei,* and *L. rhamnosus.* The lack of insertions in these species may be a direct indicator of mechanisms of genome stability. The indels found had low percentages, with insertions ranging from 0.001% - 1.5% and deletions varying between 2% to 17% (Figure 2A). The only species with higher deletion percentage was *L. plantarum,* with 17%, while other species had insertions varying between 2%-9% (Figure 2A). These findings emphasize the predominance of substitutions as the primary mode of genetic variation in the core pangenome of the *Lactobacillaceae* family.

The correlation between the variants count, and the number of core variant genes for each strain was calculated to understand the potential interplay between core genes and variability, which could shape the evolution and adaptation of these bacteria (Figure 2B). Pearson’s correlation coefficient was calculated and found to be 0.85, suggesting a strong positive correlation between the variability and core variant genes. Most strains, on average, demonstrated a median of 525 core variant genes within a range of 400-650 and a median of 1200 variants in the 800-2000 range (Figure 2B). Only a few strains from certain species exhibited greater and smaller quantities of core variant genes and variants. Notably, strains of *L. parabuchneri* and *L. ruminis* recorded the smallest and largest numbers of core variant genes and corresponding variants, respectively.

Further, the unique variants present in each core variant gene for every species were identified. For this analysis, every distinct amino acid substitution in both the consensus sequence and the variant was considered. The resulting box plot describes the distribution and variability of unique variants within core variant genes across the 26 species (Figure 2C). It was found that unique insertions within each gene of the *Lactobacillaceae* family varied from 1-16 amino acids, while deletions ranged from 1-303 amino acids, and with substitutions ranging from 2-6062 amino acids.

Very few core variant genes with a high number of unique variants were observed. Interestingly, most core variant genes encoded for hypothetical proteins in these highly variant core genes, and the others encoded for the membrane proteins, except an adhesion exoprotein (Table S2). In a case-by-case analysis, *L. plantarum* and *L. reuteri* had core variant genes with higher unique substitutions with a median count of 30 and 40, respectively, compared to other species. In the case of indels, *L. acidophilus* had core variant genes with higher unique deletions with a median count of 9, and the median count of unique insertions in all the species was 1 (Figure 2C). This analysis shows that most core variant genes exhibited a relatively low number of unique insertions and deletions and a moderate number of substitutions.

To further investigate the selection pressures acting on these species through the mutations, the average of unique variants per core variant gene in all 26 species was calculated and ranked from higher to lower values (Figure 2D). It was observed that *L. reuteri* and *L. salivarius* showed the highest unique mutations per gene, while *L. acidophilus* showed the lowest number of mutations. These results suggest that the strains in these three species are exposed to different types of pressures.

### 3.3 Purifying selection plays a dominant role in the natural sequence variation in the core pangenome of the *Lactobacillaceae* family

In the *Lactobacillaceae* family, the ratio of nonsynonymous to synonymous codon substitutions (dN/dS) for each ORF was employed to identify the intensity and kind of selection pressures affecting each core gene. Genes with dN/dS ratios greater than one are influenced by diversifying (positive) selection pressure, while dN/dS ratios less than one indicate purifying (negative) selection pressure. This means that if natural selection is promoting changes in the protein sequence, the dN/dS ratio is expected to exceed one; on the other hand, if natural selection is restraining changes in the protein sequence, the ratio is likely to be less than one (Kimura, 1977; Yang and Bielawski, 2000).

In strains of the *Lactobacillaceae* family, it was observed that purifying selection plays a dominant role in the natural sequence variation of most core genes (25,191/28,538) (Figure 3A; Figure 3B). These core genes may be conserved due to the fitness costs associated with changes, implying a high degree of functional conservation across the family.

At the species level, all except *L. helveticus* and *L. acidophilus* were found to be influenced by purifying negative selection pressures (Figure 3C). In the case of *L. acidophilus*, the core genes with ratios for positive and negative selection pressure were observed to be almost equal (Figure 3C). In *L. helveticus,* the number of core genes with dN/dS ratios less than one was higher than that of core genes dN/dS ratios greater than one, indicating diversifying positive selection pressure (Figure 3C; Figure 3D).

### 3.4 Core alelleome of *L. plantarum* strains is highly conserved

*L. plantarum*, which had 611 strains and the highest amount of amino acid variants (507,116), was further studied to understand its core alleleome. The alleleome included 1161 core genes, out of which unique substitutions were found in 1157 core genes (99.65%) (Figure S4 A) and unique deletions in 164 core genes (14%). Insertions were not observed in this species’ core genes (Figure S4 B). This distribution implies that almost all the core genes in *L. plantarum* strains have undergone substitution, while deletions are quite an infrequent phenomena in the core pangenome.

The core alleleome, encompassing the substitutions in the alleles of *L. plantarum*’s core genes, serves as a representation of core genetic diversity. Comparing naturally occurring amino acids at specific positions to their respective substitutions provides insights into amino acid variability patterns within a core gene.

The alleles of each core gene were aligned, and from this alignment, the amino acid with the highest frequency of occurrence was calculated per position. If the most frequent amino acid is termed ‘Dominant’ and the substituted amino acid ‘Variant’, the positional comparison of the total count of these amino acids (where the count represents the occurrence of a particular amino acid in the number of strains) will represent the alleleome for a single core gene. This positional occurrence of ‘Dominant’ and ‘Variant’ amino acids gives the pattern of substitutions of amino acids across the alleles of a core gene (Figure 4A). If mapped onto the protein structure, these substitutions could be used to identify the hotspots of the mutations, especially if they occur at functional sites of the proteins, such as active or binding sites (Figure 4B). To clarify further, the TrpF gene from *L. plantarum*, encoding the enzyme Phosphoribosyl anthranilate (PRAI), is involved in the third step of the Tryptophan biosynthesis pathway. It converts N-(5’-phosphoribosyl)-anthranilate to 1-carboxyphenylamino)-1-deoxyribulose-5-phosphate (CdRP) (Henn-Sax et al., 2002). The substitution pattern of TrpF amino acids showed many amino acid substitutions at different positions at the sequence level (Figure 4A) and structure level (Figure 4B). To determine this enzyme’s functional site, the consensus sequence’s binding site was predicted using the COUCH server (Yang et al., 2013). A specific amino acid substitution, cysteine (C) by tyrosine (Y), at position 7 (C7Y) was observed at the predicted binding site of the enzyme in one of the strains of *L. plantarum (*Figure S5). The amino acid ‘C’ at binding site position 7 has been reported to be necessary for the enzyme activity in Tryptophan biosynthesis in *Thermotoga maritima* (Henn-Sax et al., 2002). The mutation of this residue has been shown to make the enzyme inactive (Henn-Sax et al., 2002). This suggests that, in *L. plantarum*, the substitution in the enzyme’s binding site may alter the enzyme’s activity, which could be further confirmed with experimental studies. Thus, the alleleome identifies important mutation trends and points to functionally significant sites or structures.

Further, the substitutions in the alleles of each core gene, without considering the position, can be visualized using a histogram that displays the frequencies of both the ‘Dominants’ and ‘Variants’ (normalized to a range of 0 < c ≤ 1, relative to the total number of strains containing the core gene) (Figure 4D). Combining all the ‘Dominants’ and ‘Variants’ irrespective of position, gives us a complete *L. plantarum* core alleleome framework (Figure 4C, Figure 4D). This alleleome represents the natural sequence substitution pattern of 611 *L. plantarum* core pangenome sequences from varied niches, corresponding to 708,441 amino acid alleles across 1161 core genes containing 293,062 amino acid positions. This alleleome provides a comprehensive representation of amino acid substitutions across all the core gene sequences of the studied *L. plantarum* strains.

The conserved region of the core alleleome is the region where the dominant amino acid frequency is greater than 90% (c ≥ 0.90) (Catoiu et al., 2023) (Figure 4D). This conserved region of the core alleleome consists of 286,564 (89.96%) amino acid positions where the dominant amino acid frequency is at least 90% (c ≥ 0.90) (Figure 4D). In this conserved region, there are 275,623 amino acid positions (86.53%) for which there is a complete absence of substitutions (c = 1.0) amongst the strains (Figure 4D). Furthermore, there are 4671 positions (1.466%) for which the dominant amino acid frequency is 99% or greater (0.99 ≤ c < 1.0), while 6270 positions (1.9684%) have the dominant amino acid frequency higher than 90% (0.90 ≤ c < 0.99). These results demonstrate that the *L. plantarum* core alleleome is highly conserved.

### 3.5 *L. plantarum* core pangenome variants are mostly found in “Fermented food” and “Feces”

Further, the distribution of core gene variants across different isolation sources was investigated to identify their prevalent sources (Figure 5A). The 611 strains of *L. plantarum* were derived from 235 distinct sources. These sources were classified into 53 categories, encompassing diverse origins such as raw cow milk, plants, vegetables, fermented food, dairy products, probiotics, feces, and humans. The occurrence of core gene variants across strains was evaluated. This analysis would describe the sources rich in variants and the number of strains in which these mutations occurred.

The analysis revealed the highest occurrence of variants, with 19% found in the source ’feces’ across 132 strains, 17% in fermented food across 102 strains, and 19% associated with ’unknown sources,’ detected in 111 strains (Figure 5A). The source of fermented foods encompassed a variety of items such as kimchi, fermented cereal, kefir, silage, bamboo shoots, and others. There was also a noticeably high variant percentage in distinct sources like pickles, cheese, bakery products, and other food types. On a more general scale, the wide-ranging core gene variants in *L. plantarum* are marked by two primary sources: fermented food and feces.

Additionally, a notable percentage of variants were identified from unknown sources. The significant portion of variants from unknown sources suggests unexplored environments where *L. plantarum* may be adapting.

### 3.6 *L. plantarum* core pangenome variants are enriched in specific COG categories

In the pangenome study of *Lactobacillaceae*, specific genes were identified and assigned to COG categories (Rajput et al., 2023). These COGs were mapped with the core genes corresponding to variants of *L. plantarum*, allowing for the investigation of how the variants are distributed across distinct COG categories. The enrichment analysis of these variants within core genes was conducted to assess their distribution.

The analysis of the 19 COG categories revealed that variants were statistically enriched in 6 specific COGs within the core pangenome, namely ‘-’, ‘E’, ‘I’,‘J’, ‘K’, and ‘L’ (Figure 5B). These enriched COGs are involved in metabolic functions (categories ‘E’ and ‘I’), with the individual CGOs responsible for Amino acid transport and metabolism (‘E’), Lipid transport and metabolism (I), and Information storage and processing (categories ‘J’, ‘K’, and ‘L’), with Translation (‘J’), Transcription (‘K’) and ‘Replication, Recombination, and repair’ (‘L’), and functions still need to be characterized (category ‘-’).

### 3.7 *L. plantarum* core alleleome is largely descriptive of “conservative” and “moderately conservative” effects

The Grantham score is a metric used to evaluate the impact of amino acid substitutions based on chemical characteristics such as polarity and molecular volume (Grantham, 1974). The scores are categorized into classes reflecting increasing degrees of chemical dissimilarity: ‘conservative’ (0-50), ‘moderately conservative’ (51-100), ‘moderately radical’ (101-150), and ‘radical’ (≥151). Grantham scores for the observed substitutions were calculated and their distribution was assessed across non-synonymous codons and amino acids (Figure 5C, 5D, 5E). The anticipated effects of substitutions were investigated by analyzing these codon substitutions through their corresponding amino acids. The analysis of the core alleleome comprising 2.37 million codon substitutions attributed 80% of it (1.9 million) to synonymous substitutions, while non-synonymous substitutions accounted for the remaining 20%, or 0.47 million. A proportions Z-test yielded a p-value of 0.0 (Figure 5C), clearly demonstrating significant differences in the occurrences of these substitutions. The analysis revealed significant enrichment of synonymous substitutions among the codon substitutions, pointing to a form of genetic variation that does not lead to protein structure or function changes. Furthermore, the analysis revealed that non-synonymous substitutions (resulting in equivalent amino acid changes) had moderate Grantham score values, with few substitutions classified as ‘moderately radical’ (Grantham score 170). The global average score was 71, placing them in the ’moderately conservative’ category of substitutions (Figure 5C).

In the analysis of *L. plantarum*’s core alleleome, a substantial predominance of variants characterized by ‘conservative’ (230,401;49%) and ‘moderately conservative’ (175,281;37%) effects were observed, while variants with ‘moderately radical’ effect were found in a smaller proportion (49,267;10%). Only 4% of the variants (19,870) were responsible for ‘radical’ effects. Even though the ‘conservative’ and ‘moderately conservative’ categories will lead to amino acid modifications, the protein’s overall properties are expected not to be significantly altered. Despite their rarity, these ’radical’ variants could significantly influence protein function, particularly if they are located in active or binding sites. These results highlight that within the core alleleome of *L. plantarum*, 86% of the substitutions are unlikely to influence the structure (‘conservative’ and ‘moderately conservative’). Meanwhile, 10% might affect the structure (‘moderately radical’), and a mere 4% indicate possible structural alterations (‘radical’). Such infrequent yet potentially significant substitutions could play a key role in deciphering the organism’s distinct functions or adaptive capabilities. In summary, the core alleleome of *L. plantarum* is predominantly characterized by ‘conservative’ and ‘moderately conservative’ mutations, with very few radical variants that could significantly affect protein function.

The amino acid substitution abundance and their corresponding average Grantham scores were examined to further assess the severity of substitutions across the core alleleome (Figure 5D). Substitutions starting with amino acids G, F, M, and H were found to be predominant in the core alleleome, and they were characterized by average Grantham scores in the range of 25-100 (Figure 5E). This classifies these substitutions within the ‘conservative’ and ‘moderately conservative’ categories. Further, specific substitutions starting with amino acids C, W, and Y fell into the ‘moderately radical’ class, with average Grantham scores of 176, 117, and 117, respectively (Figure 5E). However, the observed percentages of these amino acids were minimal (C-0.6%, W-0.1%, and Y-2.5%). The rarity of substitutions with ‘moderately radical’ scores involving C, W, and Y suggests a limited contribution to functional variation within the organism’s proteins. This pattern supports the hypothesis that the majority of observed substitutions may not lead to significant alterations in overall protein functionality.

In summary, these findings contribute to the understanding that the core alleleome of *L. plantarum* is principally composed of amino acid substitutions that maintain protein functionality, falling predominantly within ‘conservative’ and ‘moderately conservative’ effects.

### 3.8 *L. plantarum* demonstrates core genetic diversity across its phylogroups, complemented by distinct phylogroup-specific variants

The *L. plantarum* core genes phylogenetic tree, constructed by aligning the core genes of 611 strains, represents the evolutionary relationship between the strains (Figure 6A). These strains were further mapped with phylogroups and isolation sources. The nine distinct phylogroups were revealed using Mash from pangenome analysis of 611 *L. plantarum* strains (Rajput et al., 2023). The phylogenetic analysis revealed that even within the same phylogroups, strains are not closely related, indicating substantial core genetic variations among these strains. It was observed that strains belonging to the same phylogroups are grouped into various subclusters within the phylogenetic tree. For example, within phylogroup I, the strains were observed to segregate into four specific subclusters. Similar trends were also observed in phylogroups ‘C’, ‘D’, and ‘E’. In the phylogenetic tree, two distinct branches were obtained, each forming clusters of strains corresponding to different phylogroups: one cluster hosted phylogroups ‘A’, ‘B’, ‘C’, ‘D’, and ‘E’, while the other primarily hosted phylogroups ‘F’, ‘G’, ‘I’, and ‘H’ (Figure 6A).

Interestingly, the analysis did not reveal a similar relation between the isolation sources and the phylogroups; instead, they appeared to be broadly dispersed throughout all phylogroups. While few isolation sources were predominant and uniquely tied to specific phylogroups, some exceptions were noted. For example, although the source “feces” was closely related to phylogroup C, it was also related to all other phylogroups (Figure 6A). A similar trend was observed with the “pickles” source with a closer relation to phylogroup B and all other phylogroups. Moreover, the sources closely related to all phylogroups included fermented food, cheese, and dairy products. This observation may signify that these particular sources host higher core genetic differences.

Further, with an emphasis on identifying phylogroup-specific unique variants, the distribution of core variant genes across these *L. plantarum* strains was evaluated. This was accomplished by constructing a presence/absence matrix representing the core variant genes. Moreover, these strains were aligned with predetermined phylogroups, enabling the grouping of all core variant genes (Figure S7). This investigation uncovered a specific cluster of core variant genes uniformly represented in all analyzed strains and their corresponding phylogroups. Moreover, it also identified core variant genes uniquely associated with individual phylogroups, highlighting the intricate core genetic relationships within the studied strains.

Interestingly, phylogroups ‘A’ and ‘C’ demonstrated distinct patterns of core variant gene sets relative to other phylogroups (Figure S7). Specifically, phylogroup ‘C’ shared a common set of core variant genes either present or absent across all the strains, with only rare exceptions. In contrast, phylogroup ‘A’ had the highest number of core variant genes present across all its strains, with only a few exclusive to specific strains (Figure S7).

In summary, the cluster map showcased two distinct patterns within the phylogroups. Firstly, specific core variant genes were prevalent across all phylogroups and were prominent in most strains. Secondly, some core variant genes appeared infrequently, acting as unique identifiers for certain strains or phylogroups.

### 3.9 Phylogroup analysis in *L. plantarum* identifies unique variants, emphasizing adaptability and industrial strain differentiation

The core variant gene distribution was further assessed to identify and analyze the core genes unique and common to specific phylogroups of *L. plantarum*. An upset plot was constructed for this analysis, visualizing the core variant genes and their corresponding phylogroups (Figure 6B). A key finding from this investigation was that 399 core variant genes (34%) were common across all nine phylogroups (Figure 6B). The analysis also noted that a comparatively smaller number of core variant genes were shared among different combinations of phylogroups, with fewer than 9 groups ranging between 6 and 53 core variant genes (0.5% - 5%), as opposed to the 399 shared core variant genes. This suggests that, although there is a considerable shared genetic foundation, each phylogroup still possesses a unique set of core genes.

It was found that certain sets of core variant genes were exclusive to specific phylogroups (Figure 6B). To illustrate further, the purple node contains five unique core variant genes specific to phylogroup ‘I’, while the orange node encompasses four unique core variant genes specific to phylogroup ‘E’. Blue, green, and red nodes also represent exclusive core variant genes specific to phylogroups ‘B’, ‘F’, and ‘G’, respectively. Five of the nine investigated phylogroups had specific, unique core variant genes. This observation underlines the existence of core genetic differentiation within *L. plantarum* phylogroups.

These five phylogroups, namely ‘B’, ‘F’, ‘I’, ‘G’, and ‘E’, were further evaluated for their variants and functional annotations. Within these phylogroups, 13 core variant genes were found to exhibit unique variants. However, three core variant genes were classified as hypothetical and subsequently excluded from further examination. The remaining core variant genes and their corresponding functions were depicted using a treemap (Figure 6C). The phylogroup ‘E’ and ‘I’ consisted of two core variant genes encoding ribosomal proteins, a result also noted in phylogroups ‘F’ and ‘G’. Among ten distinct core variant genes, six of them, each corresponding to a specific phylogroup, were encoded for ribosomal proteins. This underlines a noteworthy genetic feature shared across different phylogroups.

The variants and their positions and associated isolation sources of these phylogroup-specific core variant genes were further assessed. This was demonstrated by the variant positions and strains based on their isolation sources, with colors denoting predicted substitution impacts (Figures 7A, 7B, and Figures S8A, S8B, S8C). These variants were also depicted on 3D consensus protein structures, marked by the variant’s positions and the consensus amino acids present (Figures 7A, 7B, and Figures S8A, S8B, S8C).

In phylogroup I, the highest number of variants was detected. Notably, the unique variants across all five phylogroups were isolated from two primary sources: human feces and fermented foods. There were also a few instances where the variants were found in more rarefied sources such as ogi, cactus, raw cow milk, and unknown sources. In this context, “fermented food” encompassed several sources, including kefir, cacao bean fermentation, sour bamboo shoots, fermented soybean, kimchi, and ripening beef.

## 4. Discussion

The alleleome of core genes for 2,447 strains of the *Lactobacillaceae* family of 26 species was derived using the consensus approach to understand and explore the natural sequence diversity at the nucleotide and amino acid sequence levels. This approach reveals natural sequence and structural variations by contrasting them with the consensus sequence and structural attributes. Leveraging this approach ensures capturing variations that might be missed when using a specific strain as a reference.

Mutations within the core genes could affect their functional capability, thereby affecting key aspects of the organism such as survival, metabolic competence, stress tolerance, probiotic characteristics (Lebeer et al., 2008), and organismal fitness (Kronenberg et al., 2018). The mutation analysis indicated that insertions and deletions were infrequent, while substitutions were prevalent in the *Lactobacillaceae* family. The dominance of substitutions over deletions and insertions in the *Lactobacillaceae* family’s core alleleome indicates that these bacteria primarily undergo minor genetic changes (likely point mutations) that influence gene function, potentially further driving evolutionary trajectories and enhancing their adaptive capabilities. Insertions in the critical regions of genes are expected to cause changes in the function or expression of a gene. Therefore, these are generally maintained by various error-correction systems to restrict their disruptive functions (Patel, 2016). Notably, *L. paracasei*, *L. rhamnosus*, *L. plantarum*, *L. crispatus*, and *L. inhae* had no insertions. The lack of insertions indicates genome stability in these species, suggesting an evolutionary strategy to retain core gene integrity. These findings emphasize the predominance of substitutions as the primary mode of genetic variation in the core alleleome of the *Lactobacillaceae* family. The lower percentage of deletions and the negligible presence or absence of insertions in most species suggest a certain level of core genome functional stability through selection and error-correcting mechanisms acting against the occurrence of these types of mutations.

The examination of unique variants in each core variant gene revealed that most core variant genes in the family had few unique insertions and deletions with a moderate number of substitutions. The differences in size ranges of unique insertions, deletions, and substitutions were noted in the core variant genes of the *Lactobacillaceae* family members. This indicates a notable intraspecific diversity, which could affect strain adaptability and functionality in different environments. A limited number of core variant genes exhibited a high count of unique variants. Notably, most of these variable core genes encoded hypothetical proteins, with the remainder mostly linked to membrane proteins. These genes have often shown variations in response to specific conditions in bacteria (Ijaq et al., 2022; Rahman et al., 2022; Silhavy et al., 2010). This suggests that the observed higher variability in these core genes could indicate constant adjusting of their functionality in response to different conditions.

*L. reuteri* and *L. salivarius* showed the highest unique variants per core variant gene, while *L. acidophilus* showed the lowest unique variants per core variant gene. This result indicates that *L. reuteri* undergoes significant evolutionary changes or has higher adaptability to environmental conditions. The high mutation rate in *L. reuteri* may indicate a higher capacity for adaptability and survival under changing environmental conditions or distinct evolutionary pressures. *L. salivarius* and *L. reuteri* species are associated with diverse vertebrate host ranges and colonization sites (gut, oral cavity, vagina), indicating the vertebrate-associated lifestyle is the outcome of a long-term evolutionary process (Duar et al., 2017). *L. reuteri* strains are associated with vertebrate hosts (the human oral cavity, vagina, and intestinal tract, primates, other mammals, and birds) and *L. salivarius* with humans, rodents, birds, horses, cattle, swine, primates, and other mammals (Duar et al., 2017). These findings underscore the resilience and adaptability of these species. Conversely, *L. acidophilus*’s low mutation rate might hint at a stable ecological niche with less fluctuating selection pressures. This result has been supported by the evidence that although *L. acidophilus* has been found in various human-associated sources, recent phylogenomic analysis has determined that its most likely natural habitat is the gastrointestinal tract (GI) tract (Claesson et al., 2008). Since its primary environment is the GI tract, it might necessitate fewer variations for adaptation compared to *L. reuteri* and *L. salivarius*, which inhabit a range of sources. These results suggest that the strains in these three species are exposed to different pressures.

Further, the selection pressures acting on core genes using the dN/dS ratio for each ORF showed that purifying selection predominantly influenced the sequence variation of most core genes (25,191/28,538) in strains of the *Lactobacillaceae* family (Figure 3A and 3B). Excluding *L. helveticus and L. acidophilus*, all species demonstrated purifying negative selection pressures (Figure 3C). This highlights distinct evolutionary adaptive strategies in *L. helveticus* and *L. acidophilus*, tied to their distinct ecological roles. Where most species suppress changes, these two demonstrate increased evolutionary adaptability. Diversifying positive selection pressures are evident in *L. helveticus* (Figure 3C and 3D), while *L*. *acidophilus* displayed a near balance between positive and negative selection pressures (Figure 3C). This suggests *L. helveticus*’s adaptability might be driven by its thermophilic traits and dairy fermentation roles, potentially exhibiting constant exploration strategies for further stress resilience. This is consistent with the noted variants in its strains from sources like fermented food, fermented milk, cheese, alcoholic drinks, dairy products, bakery products, and probiotics (Figure S6). Meanwhile, *L. acidophilus*’s balanced selection strategy potentially enables it to maintain core functions and adapt, optimizing its niche role as a probiotic in the gut.

*L. helveticus* is an industrially important thermophilic starter culture in the dairy industry, especially in fermented milk and cheese production. *L. helveticus* is predominantly found in the dairy domain in cheeses and fermented milk. It includes a diverse array of products, such as koumiss (Sun et al., 2010), tarag (Liu et al., 2012), cheeses (Hebert et al., 2000), qula (Bao et al., 2012), natural whey culture (Ercolini et al., 2008), and variety of fermented milk (Shangpliang et al., 2018; Miyamoto et al., 2015). Moreover, *L. helveticus* strains are used as probiotics (Taverniti and Guglielmetti, 2012; Fontana et al., 2019; Skrzypczak et al., 2020; Toropov et al., 2020), psychobiotics (probiotics that improve gastrointestinal activity as well as anxiolytic and even antidepressant abilities) (De Oliveira et al., 2023). It can also survive environmental stresses such as high temperatures or low pH, osmotic pressure, and oxygen (Taverniti and Guglielmetti, 2012). On the other hand, *L. acidophilus* is a natural inhabitant of the human gut, and various strains are employed as probiotics (Taverniti and Guglielmetti, 2012). Additionally, of the 26 species*, L. acidophilus* and *L. helveticus* are the species with the most closed pangenome and the most open pangenome, respectively (Rajput et al., 2023), where a closed pangenome reflects the lower possibility of the addition of new gene families, while the open pangenome indicates the inclusion of new gene families in a lineage. The trend towards diversifying selection in *L. helveticus* and its open pangenome nature might be due to the diverse industrial environments it encounters and its adaptive and survival abilities.

PCA was utilized to assess the links between amino acid substitutions (consensus and variant amino acid), functions (COGs), and their predicted substitution effects (See method 2.5 and section 3S in Supplementary file). Six principal components emerged, accounting for 100% of the variability; notably, the first three PCs captured 58% of the variance (Table S1). PC1 loadings indicate varying substitution counts across species, a trend potentially due to the differences in strain count between *Lactobacillus* species (Figure S1C). PC2 loadings connected the dimensions of ‘Consensus amino acid’, ‘Variant amino acid’, ‘COG category’, and ‘Grantham score’, which are potentially intertwined in shaping the evolutionary trajectory of the *Lactobacillaceae* family. This suggests a strong coupling between the type of amino acid substitutions, the gene functions they affect, and their evolutionary consequences. However, these traits showed a moderate correlation, hinting at a link between these dimensions. It indicates a complex landscape of genetic variation and functional diversity within the *Lactobacillaceae* family, likely shaped by myriad selective pressures that may be unknown or complex. This is supported by the evidence of substantial variance by PCs, suggesting the presence of either the common mechanisms or the patterns of amino acid substitutions affecting the protein functions across the *Lactobacillaceae* family.

*L. plantarum* is recognized for its adaptability and notable features, making it a prime candidate for scientific studies. The Korean Ministry of Food and Drug Safety (MFDS) has recognized it as a probiotic bacterium for its benefits in food fermentation and promoting human health (Jung and Lee, 2020). Studies have highlighted its antibacterial capabilities, ensuring food preservation and safety (Sorrentino et al., 2013; Arena et al., 2016; Danilova et al., 2019). It also holds essential properties for manufacturing vitamins, bacteriocin, probiotics, and antifungals (Yang and Chang, 2010; Li et al., 2016; Ahn et al., 2018). In a detailed study of *L. plantarum*, which had 611 strains and the highest count of amino acid variants (507,116), the defining characteristics of its core alleleome included : 1) The *L. plantarum* alleleome is highly conserved, 2) Among analyzed isolation sources, the prevalence of variants was predominantly attributed to three categories: fermented food, feces, and unidentified origins, 3) variants are enriched in the COGs related to metabolic functions (categories E and I), information storage/processing (categories J, K, and L), and unknown functions (category -), and 4) the alleleome is predominantly characterized by ‘conservative’ and ‘moderately conservative’ mutations, with sparse radical variants potentially impacting protein function. The high amount of variants in fecal and fermented food sources suggest that these environments have a variety of selection pressures, driving *L. plantarum*’s core genetic diversification. The substantial amount of variants from unknown sources points to potential yet undiscovered environments influencing *L. plantarum* adaptation.

*L. plantarum*, due to its demonstrated probiotic properties, is considered a promising probiotic candidate (Fidanza et al., 2021). *L. plantarum* has been shown to adjust its amino acid metabolism under stress (Fidanza et al., 2021). Notably, amino acid metabolism is pivotal for sustaining cellular homeostasis and promoting resistance to external stresses across many *Lactobacillus* species (Reveron et al., 2012; Heunis et al., 2014; Šeme et al., 2015). Acid tolerance is critical for survival and adaptability, allowing strains to tolerate gastric acidity and navigate bile acid challenges in the small intestine (Fidanza et al., 2021). Many *L. plantarum* strains exhibit varying degrees of acid resistance through diverse mechanisms (Kaushik et al., 2009; Guidone et al., 2013; Hamon et al., 2014). Intriguingly, a shift in the plasma membrane fatty acid composition plays a pivotal role in *L. plantarum*’s acid tolerance (Huang et al., 2016), and many strains also exhibit bile salt hydrolase activity (Wang et al., 2021), influencing lipid metabolism and transport (Wu et al., 2022). Given the marked presence of variants in COGs associated with amino acid and lipid metabolism in *L. plantarum*’s alleleome, these variants might signal the specific survival and environmental adaptability traits of *L. plantarum* strains.

In the *L. plantarum* core alleleome, out of 2.37 million codon substitutions, 80% (1.9 million) were synonymous, while the remaining 20% (0.47 million) were non-synonymous. This highlights a pronounced preference for synonymous substitutions, pointing to a form of core genetic variation that typically does not alter protein structure or functionality. Moreover, the non-synonymous substitutions, starting with similar amino acids, predominantly fell into the ‘moderately conservative’ category with a global Grantham score average of 71. Notably, only a few had scores as high as 170, categorizing them as ‘moderately radical’ (Figure 5C). This suggests that *L. plantarum*’s core alleleome primarily consists of ‘conservative’ and ‘moderately conservative’ mutations, with limited radical variants that might significantly impact protein function. Such a genetic landscape might underpin the organism’s functional robustness, underscoring the need to delve deeper into the role of radical effect variants in *L. plantarum*’s biology. Similar findings were also observed in the alleleome of *E. coli* (Catoiu et al., 2023) and 184 bacterial species (Palsson et al., 2023); the average Grantham score of non-synonymous substitutions was found to be 67 and 63, respectively, showing that most amino acid substitutions are less severe falling in between ‘conservative’ and ‘moderately conservative’ category. Additionally, by evaluating amino acid substitution frequency and their respective average Grantham scores (Figure 5E), substitutions starting with amino acids G, F, M, and H emerged as dominant, bearing Grantham scores between 25-100 in *L. plantarum*. In a similar study on *E. coli* alleleome, substitutions beginning with L, V, and A, scoring between 40-100, were more prevalent (Catoiu et al., 2023). This illustrates the varied amino acid substitution patterns in the two organisms.

The *L. plantarum* core genes phylogenetic tree, constructed by aligning the core genes of 611 strains, represents the evolutionary relationship between the strains (Figure 6A). These strains were further mapped with phylogroups and isolation sources. In the phylogenetic tree, two distinct branches were obtained, each forming clusters of strains corresponding to different phylogroups: one cluster primarily hosted phylogroups ‘A’,‘B’,‘C’,‘D’, and ‘E’, while the other primarily hosted with phylogroups ‘F’,‘G’,‘I’, and ‘H’ (Figure 6A). This well-defined separation of strains into two primary clusters based on phylogroups may indicate unique evolutionary pathways or different selection pressures between these groups. Interestingly, the analysis did not reveal a similar relation between the isolation sources and the phylogroups; instead, they appeared to be broadly dispersed throughout all phylogroups. This dissimilarity between phylogroups and isolation source clustering might indicate that individual *L. plantarum* strains exhibit ecological versatility, adapting to diverse environmental contexts. This absence of relatedness between phylogroups and isolation sources might indicate that *L. plantarum* strains exhibit a vast ecological versatility, adapting to diverse environmental contexts. This adaptability could account for their ubiquitous presence in various habitats, such as fermented foods, dairy products, and the human gastrointestinal tract reflecting *L. plantatum*’s well known nomadic lifestyle.

The distribution of core variant genes across these *L. plantarum* strains was evaluated by constructing a presence/absence matrix of the core variant genes and mapping them with phylogroups. The analysis revealed a specific set of core variant gene in all analyzed strains and their corresponding phylogroups. The widespread nature of these shared core variant genes suggests they are crucial for essential cellular functions in the core genomes of *L. plantarum*. Interestingly, phylogroups ’A’ and ’C’ demonstrated distinct patterns of core variant gene sets relative to other phylogroups. Specifically, phylogroup ’C’ shared a set of core variant genes either present or absent across all the strains, with only rare exceptions. This suggests a mostly conserved genome in this phylogroup, perhaps indicative of specialized ecological adaptation or niche occupancy. This analysis aligns with the observations from the phylogenetic tree, showing the major niche occupancy of phylogroup ’C’ in feces and other specialized adaptations in fermented food, fermented milk, medium milk, and bakery products (Figure 6A). In contrast, phylogroup ’A’ had the highest number of core variant genes present across all its strains, with only a few exclusive to specific strains (Figure S7). This suggests that this phylogroup with high genetic diversity might inhabit a variety of isolation sources, which was evident in the varied isolation sources hosted by it (Figure 6A). This observation also aligns with the pan-genomic studies of *L. plantarum*, where this phylogroup was found to originate from a divergent set of strains, as indicated by Mash distance values, forming its distinct clade (Rajput et al., 2023; Li et al., 2022). The higher number of core variant genes in this phylogroup might indicate versatility, allowing them to strive in diverse environments.

The further evaluation of core variant gene distribution across *L. plantarum* phylogroups revealed 399 (34%) genes consistent across all nine phylogroups, emphasizing a shared genetic core crucial for primary biological functions. Despite this common foundation, unique core variant gene sets distinguished each phylogroup. Of the 9 phylogroups, 13 core variant genes stood out as unique, predominantly in phylogroups ’I’, ’E’, ’B’, ’F’, and ’G’. Of these, three were hypothetical proteins, while six encoded ribosomal proteins. Specifically, genes “DltX”, “Pts23B”, and “FabZ1” were exclusive to phylogroup ‘I’ and “CspP” to phylogroup ‘B’. Notably, the unique variants across nearly all phylogroups were isolated from two primary sources: human feces and fermented foods.

*L. plantarum*, a prevalent lactic acid bacterium in the environment, is predominantly used for the production of fermented products (Vescovo et al., 1993). During the process of production and ripening, they frequently face various environmental stresses, notably cold temperatures. The “DltX” gene encodes a protein essential for D-Ala-teichoic acid biosynthesis, contributing to the d-alanylation of teichoic acids in the bacterial cell wall. This alanylation modifies the cell wall’s charge, impacting various physiological properties (Kamar et al., 2017). This process is pivotal for gram-positive bacteria, potentially enhancing resistance to antimicrobial peptides and aiding in host cell adhesion (Collins et al., 2002). The “Pts23B” gene is a member of the phosphotransferase system (PTS) family. The PTS system plays a fundamental role in carbohydrate uptake and phosphorylation, dictating the sugars a bacterium can effectively metabolize (Rodrigo-Torres et al., 2019). “Fabz1” is commonly linked to fatty acid biosynthesis. This process is essential for bacterial cell membrane formation and has implications for other cellular functions (Zhang and Rock, 2008). A variant of this gene could suggest adaptations in membrane structure or composition, which can influence the bacterium’s interactions with its environment, stress resistance, or overall physiology. In general, the presence of these unique genes and variants in phylogroup ‘I’ suggests that this group of *L. plantarum* may have specific adaptations or characteristics related to cell wall structure, carbohydrate metabolism, and fatty acid biosynthesis. These adaptations could influence their probiotic properties, interactions with the host, and survivability in various environments.

Typically, *L. plantarum* encounters severe environmental conditions during processing and storage, which can involve freeze-drying or standard freezing. Fermented foods predominantly rely on cold for preservation. The “CspP” gene encodes cold shock proteins upregulated in response to sudden temperature decreases in *L. plantarum* (Derzelle et al., 2002). Resistance to cold stress is pivotal for probiotic attributes (Song et al., 2014). The unique variants of “CspP” in a specific phylogroup of *L. plantarum* might indicate specialized adaptations related to temperature shifts or potentially other stressors. It suggests that this phylogroup might have evolved mechanisms to handle such conditions more effectively than others.

In summary, the unique variant genes specific to phylogroup ‘I’ suggest that the strains with these variant genes (“DltX”, “FabZ1”, “Pts23B”) might possess more competitive adaptability to environments than the other strains and could possess specialized abilities related to probiotic properties, host interactions, and survivability in different environments. Phylogroup ‘B’ specific unique variant gene suggests that the corresponding strains might have evolved to adapt to the conservation system essential for fermented food preservation and other stresses pertaining to the gut environment and fermentation process. They might possess the characteristics of specialized thermal adaptations and stress tolerance.

## 5. Conclusion

The pan-genome provides a comprehensive genetic overview of a species, highlighting potential lifestyles, whereas the core genome represents universally shared genes, underscoring key biological functions and traits. The alleleome offers further in-depth exploration and analysis of sequence diversity within the genes of these species. Using the genetic consensus sequences, the alleleome aids in understanding feasible variability in gene ORFs. This could be a valuable approach for exploring 3D protein structure alterations and advancements in protein design and engineering. The current study revealed that the core alleleome of the *Lactobacillaceae* family mainly had substitutions, indicating minor genetic changes that might affect gene functions and influence evolution and adaptability. Few species hosted deletions or insertions, suggesting at genome stability and robust error correction mechanisms for the core genome in the family. Purifying selection predominantly influenced the sequence variation of most core genes in strains of the *Lactobacillaceae* family. Distinct evolutionary traits were noted in *L. helveticus* and *L. acidophilus. L. helveticus* displayed diversifying selection suggesting the adaptability emerging due to its thermophilic nature and exploration strategies for stress resilience in different environments. *L. acidophilus* displayed balanced selection pressures potentially enabling it to maintain function and optimize its unique role.

In a detailed study of *L. plantarum*, the defining characteristics of its core alleleome included: 1) The *L. plantarum* alleleome is highly conserved, 2) Among analyzed isolation sources, the prevalence of variants was predominantly attributed to three categories: fermented food, feces, and unidentified origins, 3) Core gene variants are enriched in the COGs related to metabolic functions, information storage/processing, and unknown functions, and 4) the alleleome is predominantly characterized by “conservative” and “moderately conservative” mutations, with very few radical variants potentially impacting protein function. Phylogroup-specific core variant gene analysis identified unique variants in phylogroups ‘I’ and ‘B’. These variants hint at enhanced adaptability, probiotic traits, host interactions, environmental survivability, thermal resistance, and stress tolerance of select strains. Unique variants were found in five of nine phylogroups, indicating core genetic diversity across phylogroups. These findings pave the way for understanding phenotypic differences in *L. plantarum* strains. The distinct variants offer a genomic identifier for each phylogroup, aiding in precise strain tracking, identification, and evolution studies in industrial settings.

## Acknowledgment

We thank Akanksha Rajput for providing metadata for the 26 species of the *Lactobacillaceae* family.

## Funding

This work was supported by the Novo Nordisk Foundation through the Center for Biosustainability at the Technical University of Denmark (NNF Grant Number NNF20CC0035580).

## Conflict of interest statement

None declared.

## Notes

https://github.com/anpanche/Lacto_alleleome

### Competing Interest Statement

The authors have declared no competing interest.

