## Supplementary material for "Genomic insights into *Lactobacillaceae*: Analyzing the “Alleleome” of core pangenomes for enhanced understanding of strain diversity and revealing Phylogroup-specific unique variants": Supplimentary file

### 3S Interplay between amino acid variations, functional impacts, and predicted substitution effects in the *Lactobacillaceae* family

The distribution of variants (substitution, indels) across the Clusters of Orthologous Groups (COG) categories was further investigated. COGs enable the systematic assigning of functions to genes (Koonin et al., 1998). This assignment would give us information on the distribution of variations across the critical functions in the family. The COG assignments from pangenome analysis (Rajput et al., 2023) were mapped with the core genes hosting variants across 26 species, and the percentage distribution of all variant types was plotted using a heatmap (Figure S2). The COG categories use symbols ranging from one to three letters. A single-letter COG symbol represents genes involved in one function, while two and three letters indicate genes associated with two and three COG functions, respectively. The heatmap demonstrated that specific COG categories and their combinations had abundant substitutions with little deletions and almost no insertions. High substitutions in specific COGs suggest these core genes might be undergoing adaptive evolution, where beneficial mutations are being explored. If the relationship between these substitutions, COGs, and their predicted substitution effects were understood, it would enable the association of these substitutions with their specific functions and predicted substitution effects.

One of the approaches to predict the impact of the substitutions are Grantham scores (Grantham, 1974). The Grantham scores of the amino acid substitutions were calculated for all substitutions of 26 species. PCA was performed to investigate the relationship between the amino acid substitutions with functions (COGs) and their predicted mutation effects. Uncovering these relationships could aid our understanding of the selective pressures shaping the genomes of these organisms.

PCA was carried out on the dataset using the substitution count, consensus amino acid, variant amino acid, COG category, Grantham score, and species name. PCA derived six principal components (PC), explaining 100 % variability ("Cumulative" row) within our dataset (Table S1). The first two PCs accounted for 21% and 19% of the total variance ("Proportion" row), respectively, explaining 40% of the variation, while the first three PCs explained 58% of the variance.

The variables observed with higher weights in PC1 were 'Substitution count' and 'Species' (Table S1). In PC2, the variables with higher weights were found to be 'Consensus amino acid', 'Variant amino acid' along with the 'COG category', and 'Grantham score' (Table S1). In the PCA biplots (Figure S3-1), clustering of samples was observed when the components were colored with either consensus amino acids (Figure S3-1B) or variant amino acids (Figure 4C) and with predicted substitution effects (Figure S3-1D). The PCA biplot was also colored by COGs (Figure S3-2 A), species (Figure S3 B), and amino acid substitution pairs (Figure S3-2 C). This observed clustering across all biplots suggests trends between each pair of dimensions (components representing variants, COGs, Grantham score, and species names).

*Table S1: The PCs and their correlation coefficient variables. The table also includes the explained and cumulative variance for 6 principal components.*

| | PC1 | PC2 | PC3 | PC4 | PC5 | PC6 |
|---|---|---|---|---|---|---|
| **Grantham Score** | 0.10 | 0.24 | 0.31 | 0.21 | 0.02 | 0.13 |
| **Substitution Count** | 0.67 | 0.02 | 0.00 | 0.02 | 0.06 | 0.23 |
| **Consensus Amino Acid** | 0.02 | 0.23 | 0.34 | 0.07 | 0.31 | 0.03 |
| **Variant Amino Acid** | 0.04 | 0.36 | 0.16 | 0.03 | 0.39 | 0.02 |
| **COG Category** | 0.09 | 0.31 | 0.25 | 0.21 | 0.01 | 0.14 |
| **Species** | 0.33 | 0.01 | 0.03 | 0.45 | 0.09 | 0.09 |
| **Eigen Value** | 1.26 | 1.16 | 1.09 | 0.98 | 0.87 | 0.64 |
| **Proportion** | 20.94 | 19.38 | 18.24 | 16.40 | 14.44 | 10.60 |
| **Cumulative** | 20.94 | 40.32 | 58.56 | 74.96 | 89.40 | 100.00 |

When the variables with high loading scores in PC1 (Figure S3-1A) were analyzed, it was determined that the substitution count and the species name variable loadings were in the same direction, indicating that the component captures the variations in substitution counts and species names. These two dimensions are therefore predicted as related, with the substitution count observed to contribute more than the species name (arrow lengths) (Figure S3-1A). This result suggested that the substitution count differs in the species. This relation is evident based on the strain count differences in the *Lactobacillus* species (Figure S1 C).

The PC2 showed higher loadings for all the variables except those in PC1 including 'Consensus amino acid', 'Variant amino acid', 'COG category', and 'Grantham score'. All these loadings were in the same direction, and these variables also had high weights in the other principal components, meaning that the variations in the substitutions were correlated with the COGs and Grantham scores. However, the contribution of these traits was found to be low compared to the sum of loadings, which is one. Therefore, it cannot be concluded that these traits are strongly correlated.

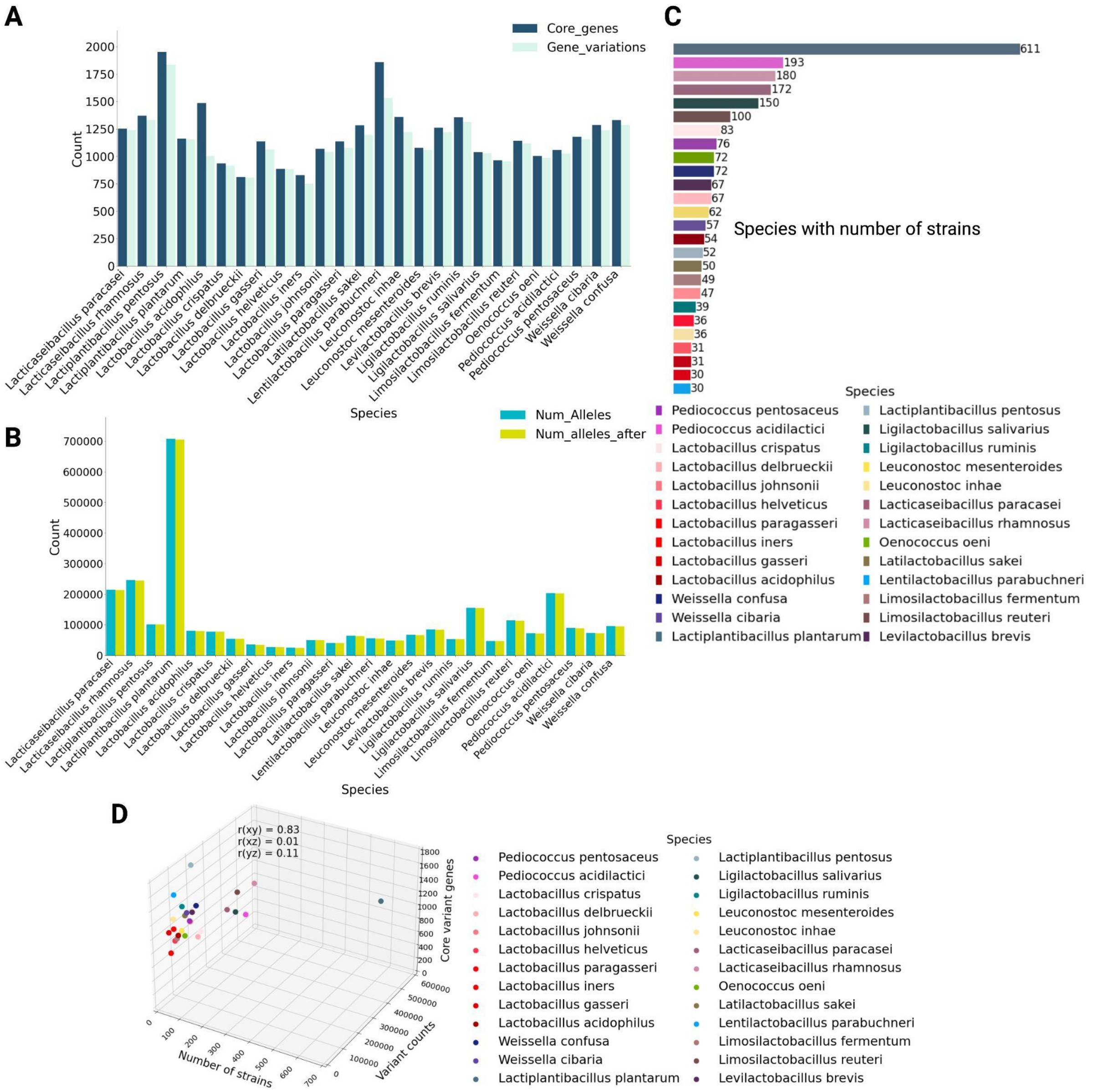


*Figure S1: A) The total number of Core genes and the number of genes with variants in each species; B) The total number of alleles before and after quality control in each species; C) The Total number of strains in 26 species; D) 3D scatter plot with correlation coefficients between the number of core variant genes with their variants count and the number of strains in each species.*

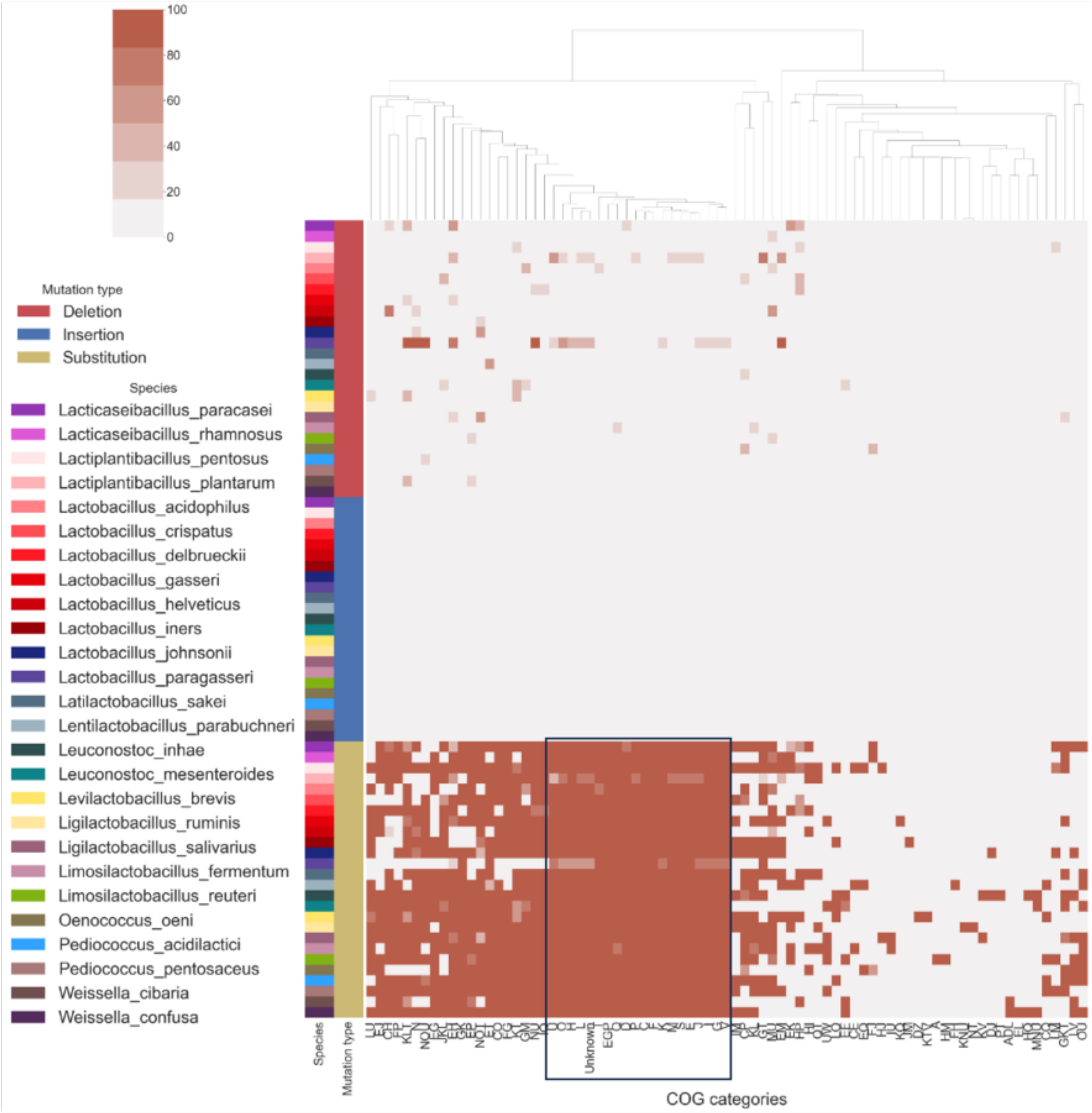


*Figure S2: Distribution of variants with mutation types (substitution, insertion, and deletion) across specific and combined COG categories in the core pangenome of each species of the Lactobacillaceae family. Each column displays the cumulative count of variants, categorized by mutation type, and indicates their occurrence in specific species within the designated COG category.*

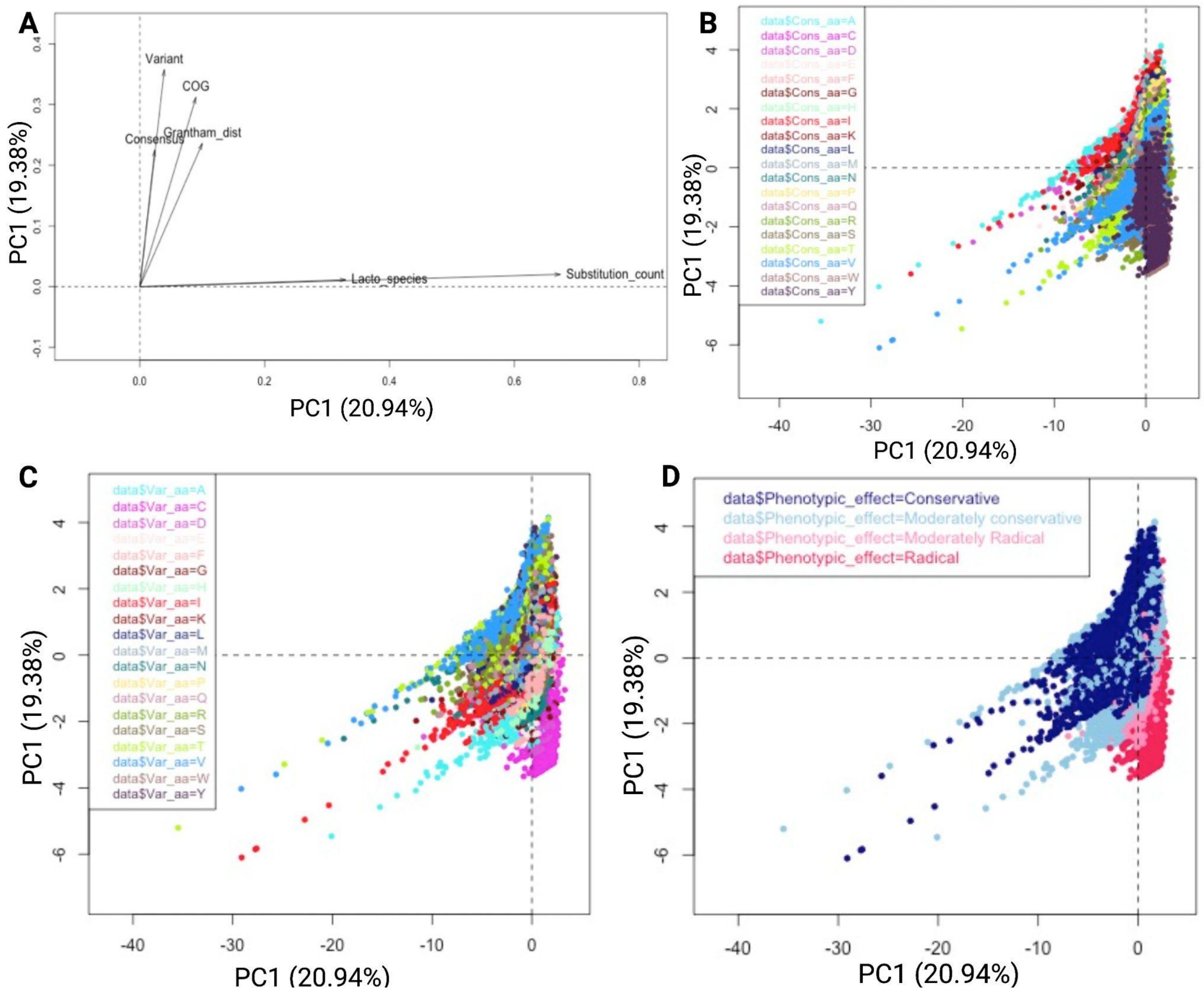


*Figure S3-1: A) Square loading of all the variables across PC1 and PC2; B) The distribution of observations of PC1 and PC2 colored by Consensus amino acids; C) The distribution of observations of PC1 and PC2 colored by Variant amino acids; D) The distribution of observations of PC1 and PC2 colored by amino acid substitution effects given by Grantham score; PCA was carried out using the PCAmixdata package in R.*

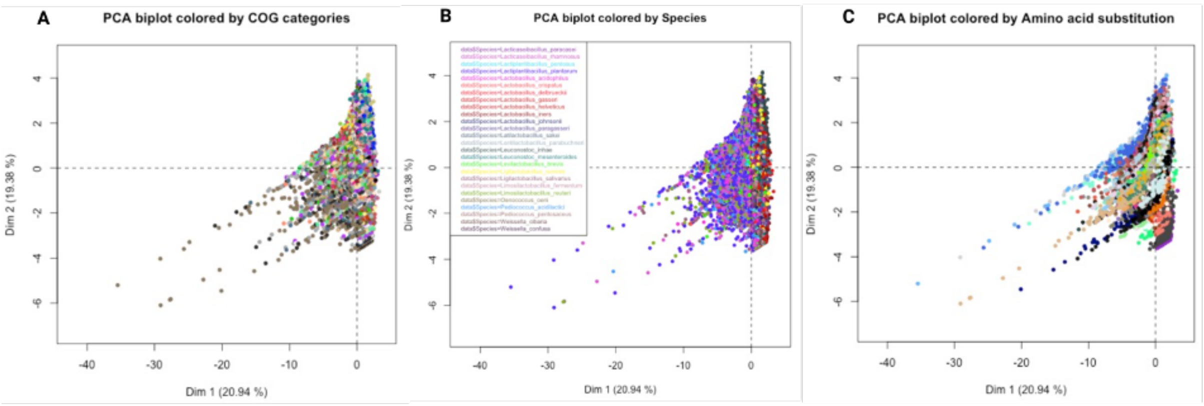


*Figure S3-2: A) The distribution of observations of PC1 and PC2 colored by COG categories B) The distribution of observations of PC1 and PC2 colored by species; C) The distribution of observations of PC1 and PC2 colored by amino acid substitution pairs.*

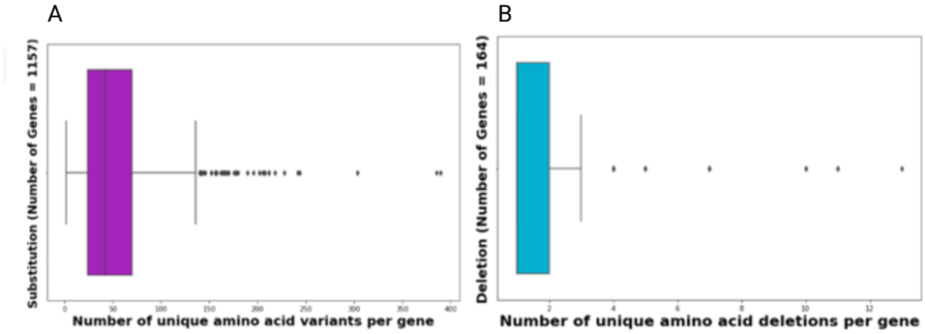


*Figure S4: General distribution of variants and genes with unique variants. The substitutions and deletions percentage ratio in L. plantarum is shown. Core genes with their unique variant count with substitution and deletion are shown. No insertion was observed in L. plantarum 611 strains.*

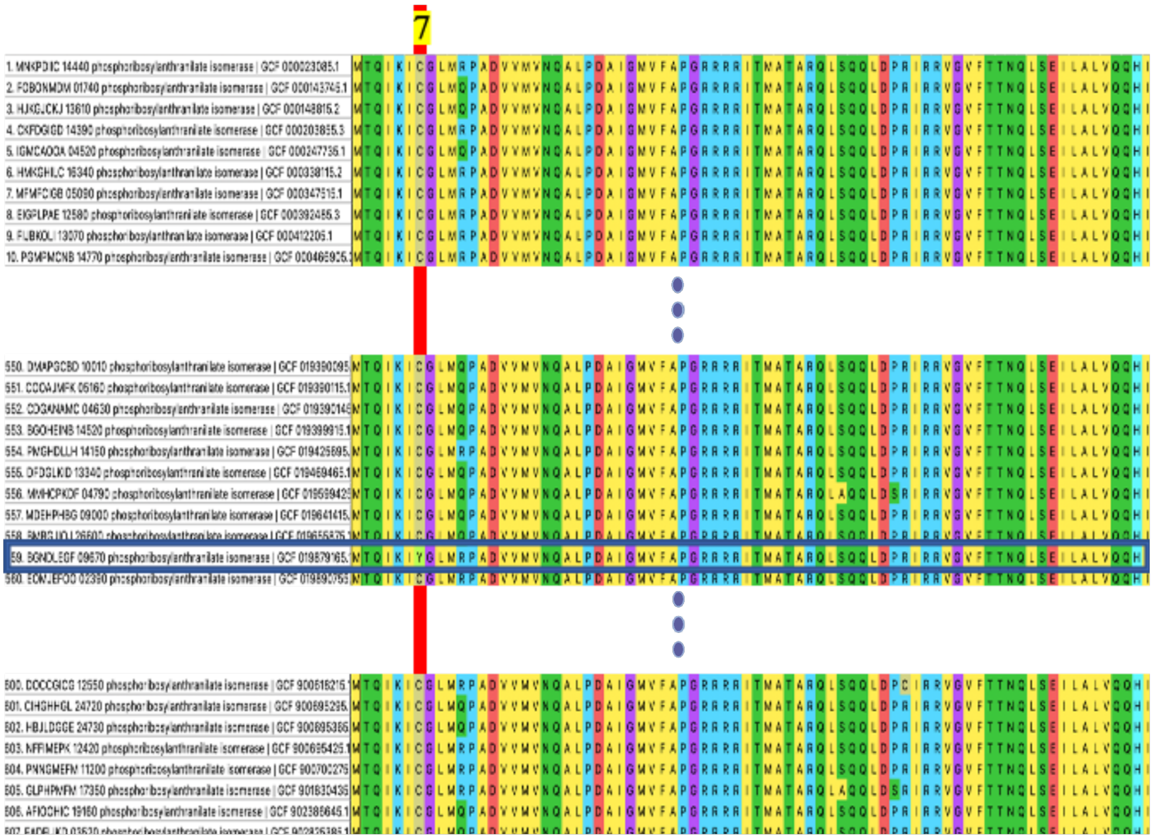


*Figure S5: Multiple sequence alignment of alleles of TrpF gene in L.plantarum strains. Position 7 highlights the occurrence of amino acid substitution patterns of cysteine (C) and Tyrosine (Y). The specific substitution C7Y was observed in only one strain of L. plantarum.*

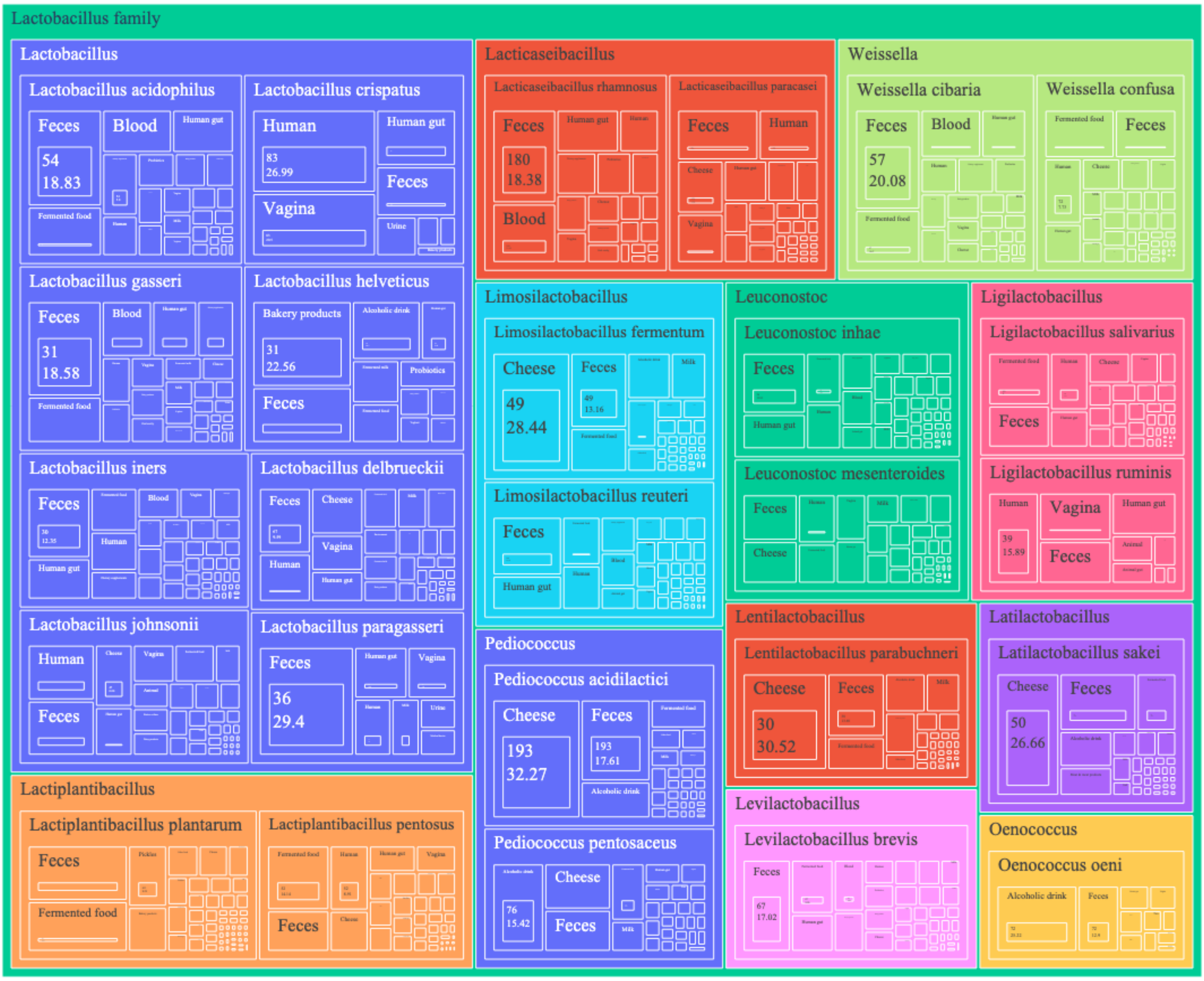


*Figure S6: Treemap of all species showing the number of strains in which the variants were observed. The squares are colored by the genera of the species. The size of the squares indicates the count of the strains with variant percentages.*

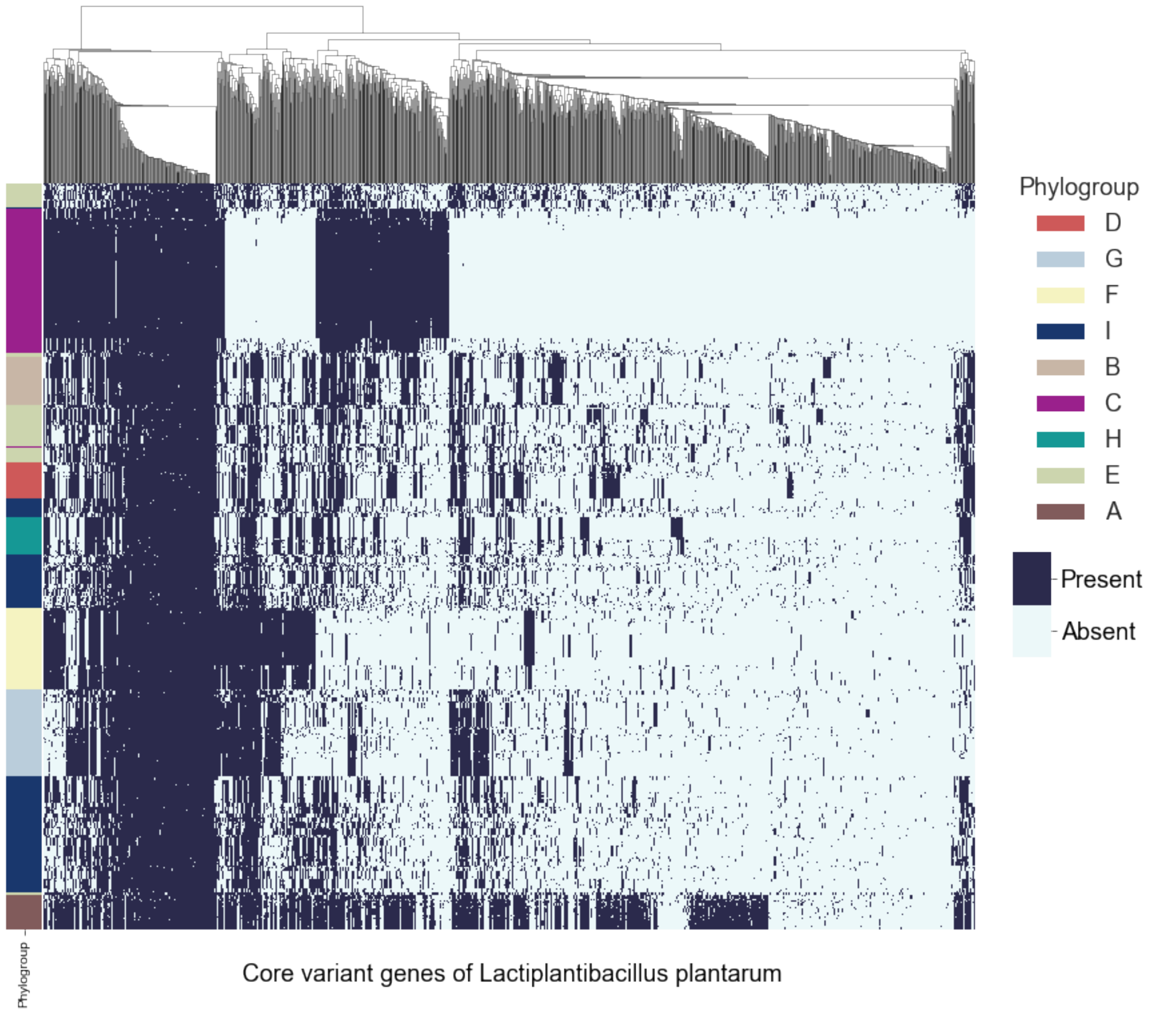


*Figure S7: Presence and absence matrix of core variant genes across phylogroups in L. plantarum.*

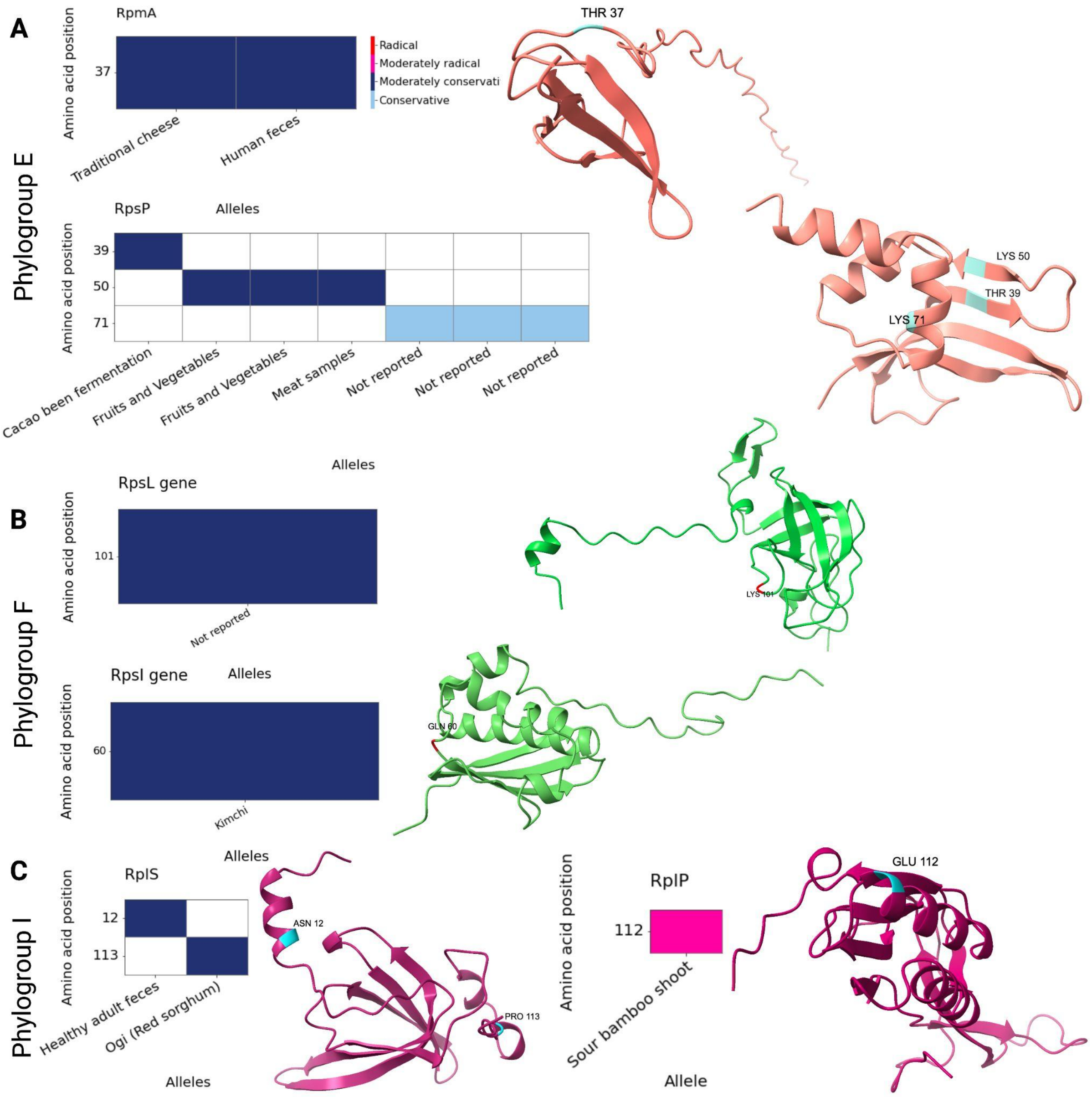


*Figure S8: A) Phylogroup E specific ribosomal protein variants of genes "RpmA", and "RpsP" with their amino acid mutation position, predicted substitution effects, and isolation sources; B) Phylogroup F specific ribosomal protein variants of genes "RpsL", and "RpsI" with their amino acid mutation positions, substitution effects, and isolation sources; C) Phylogroup I specific ribosomal protein variants of gene "RplS" with their amino acid mutation position, substitution effects, and isolation sources. The mutation positions are colored by their Grantham scores predicted effects for the genes.*

*Table S2: Core variant genes with a higher number of unique variants in the Lactobacillaceae family members, along with their encoding proteins, unique variants count, and mutation type.*

| Sr. No. | Gene | **Unique variants** | **Annotation** | **Mutation** | **Species** |
|---|---|---|---|---|---|
| 1 | group_41 | 6062 | Hypothetical protein | Substitution | *Weissella confusa* |
| 2 | group_57 | 5940 | Hypothetical protein | | *Lactobacillus iners* |
| 3 | inlI | 3264 | Adhesion exoprotein | | *Ligilactobacillus salivarius* |
| 4 | group_106 | 3190 | Hypothetical protein | | *Lactobacillus acidophilus* |
| 5 | smc_1 | 1754 | Membrane protein | | *Lactobacillus delbrueckii* |
| 6 | group_1033 | 1614 | Putative membrane protein | | *Lactobacillus johnsonii* |
| 7 | group_198 | 1078 | Putative membrane protein | | *Lactobacillus paragasseri* |
| 8 | group_122 | 1028 | Putative membrane protein | | *Lactobacillus gasseri* |
| 9 | group_47 | 1000 | Hypothetical protein | | *Lentilactobacillus parabuchneri* |
